# Distinct prefrontal top-down circuits differentially modulate sensorimotor behavior

**DOI:** 10.1101/307009

**Authors:** Rafiq Huda, Grayson O. Sipe, Vincent Breton-Provencher, K. Guadalupe Cruz, Gerald N. Pho, Elie Adam, Liadan M. Gunter, Austin Sullins, Ian R. Wickersham, Mriganka Sur

## Abstract

Sensorimotor behaviors require processing of behaviorally relevant sensory cues and the ability to select appropriate responses from a vast behavioral repertoire. Top-down modulation by the prefrontal cortex (PFC) is thought to be key for both processes but the precise role of specific circuits remains unclear. We examined the sensorimotor function of anatomically distinct outputs from a subdivision of the mouse PFC, the anterior cingulate cortex (ACC). Using a visually guided two-choice behavioral paradigm with multiple cue-response mappings, we dissociated the sensory and motor response components of sensorimotor control. Projection-specific two-photon calcium imaging and optogenetic manipulations show that ACC outputs to the superior colliculus, a key midbrain structure for response selection, principally coordinate specific motor responses. Importantly, ACC outputs exert top-down control by reducing the innate response bias of the superior colliculus. In contrast, ACC outputs to the visual cortex facilitate sensory processing of visual cues. Our results ascribe motor and sensory roles to ACC projections to the superior colliculus and the visual cortex and demonstrate for the first time a circuit motif for PFC function wherein anatomically non-overlapping output pathways coordinate complementary but distinct aspects of visual sensorimotor behavior.

## Introduction

The behavioral repertoire of animals is highly enriched by their ability to learn how to respond to sensory cues to achieve goals such as reward^1–10^. Though seemingly simple, goal-oriented sensorimotor behaviors require coordination of multiple processes. Animals receive a deluge of environmental information at any given moment and can express a wide range of motor behaviors. Hence, sensorimotor control requires attentional mechanisms that prioritize processing of relevant sensory cues and select task-appropriate responses. Studies over the past decades have identified the prefrontal cortex (PFC) as a crucial nexus for coordinating sensorimotor behaviors. Specifically, the PFC is thought to generate top-down signals that facilitate task-specific processing^11–18^. However, a fundamental outstanding question is how the anatomical organization of inputs to and outputs from the PFC enables its proposed role in sensorimotor control.

Previous work using electrical stimulation demonstrates that the same PFC area can both enhance the representation of cortical visual signals, a neurophysiological hallmark of visual attention, as well as facilitate motor responses^14, 19–21^. However, electrical stimulation is non-selective and hence these studies do not address whether specific PFC cell populations underlie sensory and motor functions. At the same time, other work suggests that the activity of functionally distinct populations of PFC neurons correlates with distinct components of sensorimotor behavior^16, 22^. An intriguing possibility is that sensory and motor functions are subserved by distinct PFC subpopulations that target specific downstream structures.

Recent studies have identified a PFC area in mice, the anterior cingulate cortex (ACC), which is functionally and anatomically poised to exert top-down control over visually guided behaviors. The ACC receives inputs from the visual cortex^23^ and exhibits visual responses at single-neuron and network levels^24, 25^. Studies employing causal manipulations using chemogenetics or optogenetics show that ACC activity is important for optimal performance on visually guided tasks^18, 26–28^. The ACC provides top-down outputs to the visual cortex (VC) and motor-related layers of the superior colliculus (SC)^27–31^, a crucial midbrain structure for response selection and other functions^9, 32–40^. Importantly, these outputs originate from non-overlapping populations of ACC projection neurons^29^, raising the possibility that these output pathways differentially modulate sensorimotor behavior. Surprisingly, optogenetic activation of ACC outputs to the VC or SC enhances the gain of stimulus-driven responses in the VC^27, 30^, suggesting a role for both pathways in sensory processing. While the function of ACC outputs to the VC in modulating cortical visual processing is consistent with previous work^14, 16^, a similar role for ACC outputs to the SC is harder to reconcile. The intermediate and deep layers of the SC are known to modulate specific motor functions and sensorimotor responses^9, 32, 36, 41–44^. Furthermore, pharmacological inactivation of the SC produces a strong deficit in visual attention without perturbing the associated modulation of visual cortical activity^45^. Thus, the precise contribution of ACC projections to the VC and SC in mediating sensory processing and motor responses remains unclear.

Elucidating the function of ACC outputs to the VC and SC in sensorimotor control requires probing the underlying visual processing and motor response components. Here, we establish a two-choice behavioral paradigm that allows us to distinguish these processes by testing mice on different cue-response contingencies. Comparing behavioral deficits induced by projection-specific inactivation of ACC output pathways across task contingencies shows that ACC projections to the SC modulate specific motor responses, while projections to the VC contribute to sensory processing. Remarkably, we find that ACC outputs to the SC facilitate motor responses by reducing the innate response bias of the SC. By using two-photon calcium imaging of ACC inputs/outputs, virus-mediated anatomical tracing, and optogenetics, we delineate specific circuit mechanisms mediating this novel effect.

## Results

### The caudal ACC is anatomically positioned to contribute to visual sensorimotor behavior

The ACC is a midline structure that spans a large extent across the caudo-rostral axis^46^. Although recent studies have implicated the mouse ACC in visual behaviors^18, 26–28, 30^, this PFC region is also associated with other diverse behavioral functions^47–49^. We used rabies virus-mediated anatomical tracing and two-photon calcium imaging to determine if the ACC contains a subregion specialized for visual sensorimotor behaviors. We identified sources of visual inputs to the ACC by injecting rabies viruses encoding GFP and tdTomato into caudal and rostral ACC, respectively (Figure 1A). Although both compartments received inputs from medial higher visual cortex, corresponding to functionally-defined anteromedial and posteromedial areas^50^, the caudal ACC also received inputs from the primary visual cortex (Figure 1B-C). Moreover, each compartment received prominent inputs from its contralateral hemisphere (Figure 1D). This anatomical organization suggests that: a) the VC provides visual information to the ACC; b) visual information is integrated in ACC activity; and c) visual information is exchanged between the two ACC hemispheres via callosal projections. We tested these predictions using two-photon calcium imaging of VC and callosal axons in the caudal ACC. We unilaterally injected GCaMP6s in the VC or the ACC, and placed a chronic cranial window over the caudal ACC to image visually evoked activity of axons (Figure 1E). While VC axons responded to stimuli presented in the contralateral visual field (relative to the site of recording), callosal axons responded preferentially to ipsilateral stimuli (Figures 1F, G; Supplementary Figure 1). Importantly, our callosal axon recordings establish that ACC neurons are visually responsive even in naïve mice, and relay visual information to the opposite hemisphere. Hence, VC and callosal inputs provide information about contra- and ipsilateral visual fields to the ACC, respectively.

**Figure 1.**
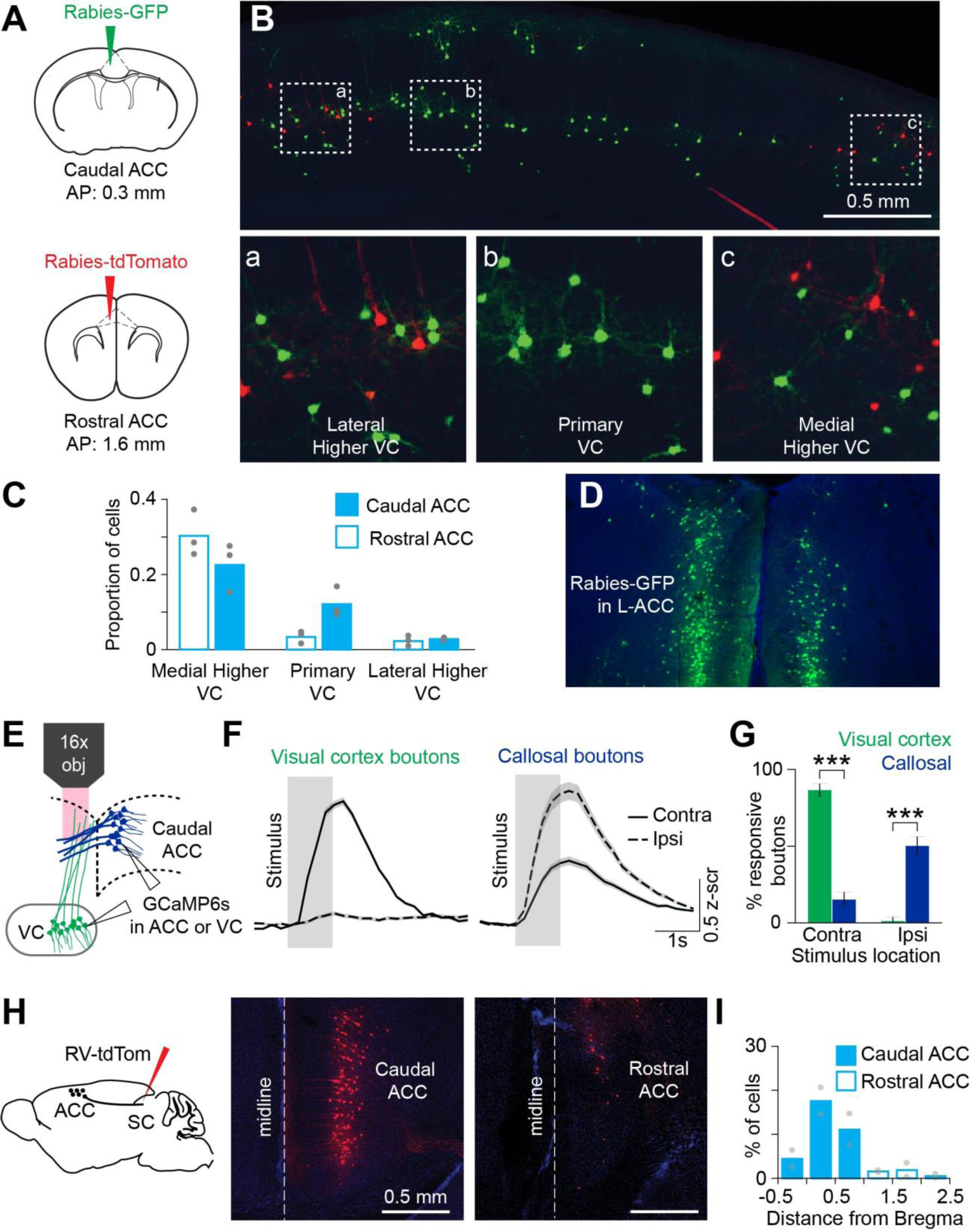
Anatomical and functional characterization of inputs to and outputs from the ACC. (**A**) Rabies viruses encoding GFP or tdTomato were injected into caudal or rostral ACC, respectively. (**B**) Back-labeled neurons in the visual cortex. Neurons in lateral higher VC, primary VC, or medial higher VC are denoted by dotted squares (a-c) on top and shown at higher magnification on the bottom. (**C**) Proportion of back-labeled neurons in various VC subdivisions projecting to caudal (solid) or rostral (unfilled) ACC (n = 3 mice). (**D**) Back-labeled callosal caudal ACC neurons. (**E**) Experimental setup for two-photon imaging of GCaMP6s-expressing visual cortex or callosal axons in the ACC while head-fixed mice passively viewed visual stimuli (black square or grating, ∼20°) presented in either hemifield. (**F**) Population-averaged responses of visually-driven VC (n = 268 boutons from 5 mice) and callosal (n = 309 boutons from 4 mice) boutons to stimuli presented in contra- or ipsilateral hemifields. Solid or dashed line is the mean response and shading shows the standard error of the mean. (**G**) Percent of visually-driven VC and callosal boutons with preferential responses to contra or ipsi stimuli. ***p < 10^-5^ (Fisher’s exact test, two-sided). Errors bars are 95% binomial confidence intervals. (**H**) Rabies viruses encoding tdTomato were injected in the SC, leading to labeling in the ACC. **I**) Proportion of back-labeled SC-projecting neurons located along the caudo-rostral axis of the ACC (n = 2 animals).

ACC projection neurons are known to provide top-down outputs to the SC^29, 30^. We used anatomical tracing to determine if these projection neurons localize to the same subdivision that receives VC inputs. Injection of a rabies virus encoding tdTomato in the SC showed a high density of back-labeled neurons in the ACC (Figure 1H). SC-projecting ACC neurons (ACC-SC) were located predominantly in the caudal subdivision (Figure 1I). This labeling was observed ipsilateral to the site of injection, establishing the unilateral nature of this projection pathway. Together, our anatomical and functional studies show that the caudal ACC integrates visual inputs and provides outputs to the SC, which could allow this PFC area to contribute to visual sensorimotor control.

### A sensorimotor behavioral paradigm for studying sensory processing and motor responses

We tested the role of the caudal ACC in visuomotor control by designing a behavioral paradigm for head-fixed mice inspired by previous work^6, 10^. This paradigm assesses sensory processing by requiring mice to detect lateralized (i.e., left or right) visual stimuli. Mice report the spatial location of visual cues by rotating a trackball in one of two directions, additionally allowing us to study motor responses. We define visual cues as contralateral (contra) or ipsilateral (ipsi) based on the side of brain hemisphere under study, which is assumed to be the left side for all figures and text (see Table 1 for targeting details and behavioral performance of mice in specific experiments). We began these experiments with the ‘inward’ cue-response contingency, in which mice move the presented stimulus to the center of the screen. In later experiments, we trained mice on an ‘outward’ contingency, in which mice move cues to the outside of the screen. For the ‘inward’ task, mice respond to ipsi cues with clockwise actions and to contra cues with counterclockwise actions (Figure 2A). We used a reaction time task design that allowed mice to make a response as soon as they could after stimulus onset. This minimized potential confounds of short-term memory and extensive movement planning associated with delay tasks^51^. Experienced mice performed well on this task (∼80% accuracy), with few timeouts (i.e., incomplete trials in which the ball is not moved to response threshold; note that timeout trials are excluded for quantifying accuracy) and responses with short latencies (∼400ms) after stimulus onset (Figure 2B).

**Figure 2.**
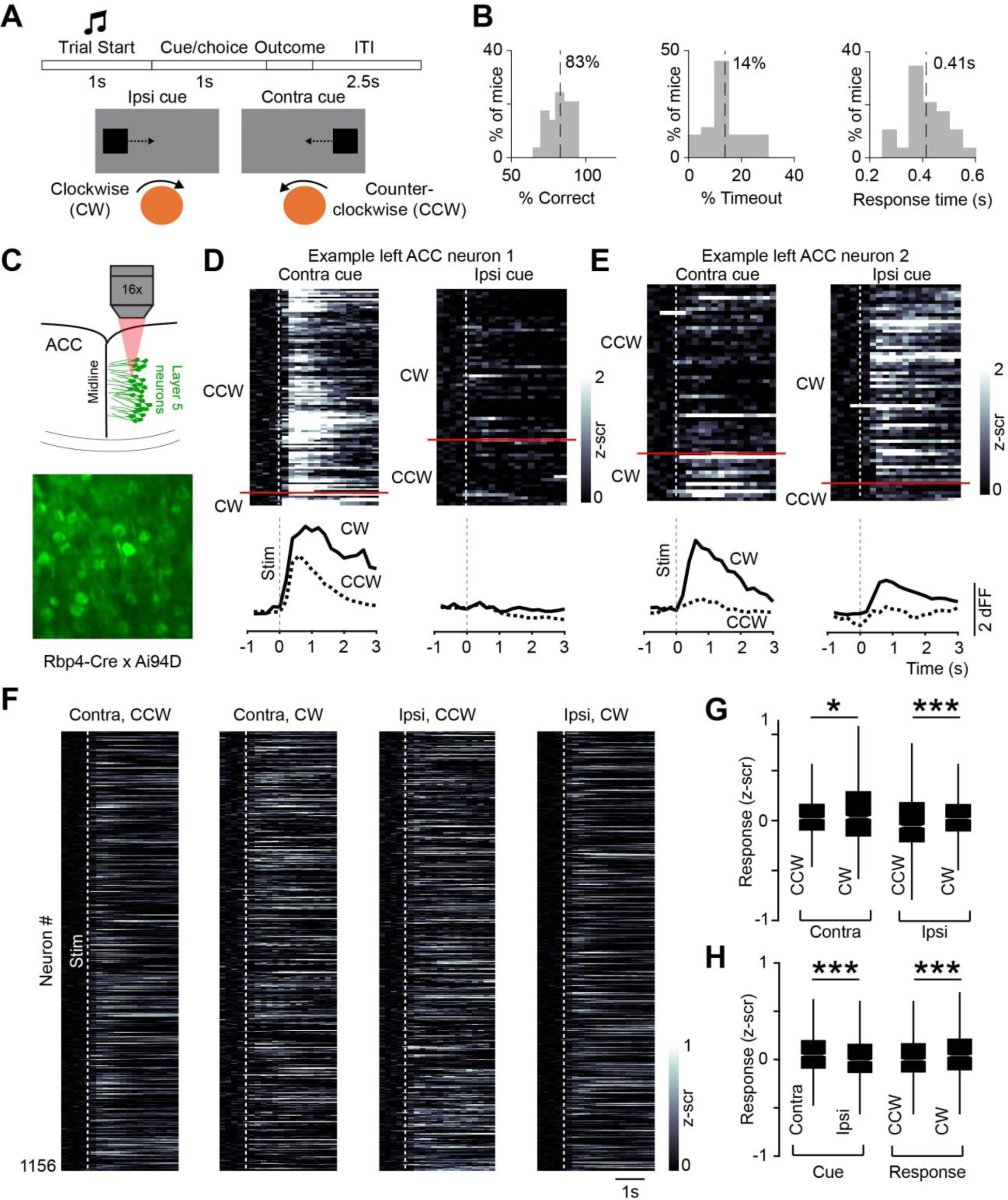
Responses of ACC neurons in a visual sensorimotor task. (A) Trial events during the task (top) and schematic showing the sensorimotor contingency Cues are labeled ipsi and contra relative to the brain hemisphere targeted for on/recording (assumed to be the left hemisphere in all figures and the text; see Table 1 for targeting details of specific experiments). Mice respond to ipsi and contra cues by rotating a trackball clockwise and counterclockwise, respectively. Counterclockwise action is the correct response for the contra cue but an incorrect response for the ipsi cue (and vice versa for clockwise action). Correct responding moves the cue to the center of the screen, whereas incorrect responding moves it to the side. (**B**) Behavioral performance of mice trained on this task for various optogenetic inactivation experiments (n = 29 mice). Dashed lines indicate mean. (**C**) Two-photon calcium imaging of GCaMP6s-expressing layer 5 neurons. Activity of 1156 neurons is reported from 5 expert mice (8 behavioral sessions, 1-3 from each mouse). Dotted lines show stimulus onset. (**D, E**) Activity of two example ACC neurons during the task. Responses on contra and ipsi cue trials with counterclockwise (CCW) or clockwise (CW) actions are shown. Rows correspond to trials. (**F**) Session-averaged activity of individual ACC neurons for the indicated trial type is shown. Each row corresponds to the same neuron across the four plots. (**G**) Average responses on contra and ipsi cue trials with counterclockwise or clockwise responses (contra, CCW vs. CW: p = 0.033, z = -2.1322; ipsi, CCW vs. CW: p = 2.87 × 10^-7^, z = -5.132). *p < 0.05, ***p < 0.005, Wilcoxon signed-rank test. (**H**) Average responses on contra vs. ipsi cue trials (p = 9.2 × 10^-9^, z = 5.745) and CCW vs. CW trials (p = 8.3 × 10^-5^, z = -3.940). ***p < 0.005, Wilcoxon signed-rank test.

**Table 1.** Performance metrics of mice included in this study.

| Mouse # | Experiment | Figure<sup>a</sup> | Implant/<br>imaging site | Behavioral performance |  |  |  |
| --- | --- | --- | --- | --- | --- | --- | --- |
|  |  |  |  | Control accuracy<sup>b</sup> | Non-laser accuracy | Timeout | Response Time (s) |
| 61 | Light Ctrl | F4-S1 | Left | 97% | 95% | 8% | 0.35 |
| 62 | Light Ctrl | F4-S1 | Left | 87% | 93% | 16% | 0.5 |
| 72 | Light Ctrl | F4-S1 | Left | 93% | 87% | 28% | 0.39 |
| 100 | Light Ctrl | F4-S1 | Left | 98% | 95% | 12% | 0.32 |
| 120* | Light Ctrl | F4-S1 | Right | 78% | 78% | 32% | 0.53 |
| 122 | Light Ctrl | F4-S1 | Left | 93% | 94% | 10% | 0.57 |
| 15 | SC inh. | F4 | Left | 65% | 61% | 4% | 0.44 |
| 68 | SC inh. | F4 | Left | 76% | 76% | 12% | 0.46 |
| 120* | SC inh. | F4 | Left | 76% | 62% | 20% | 0.48 |
| 125 | SC inh. | F4 | Left | 77% | 67% | 14% | 0.36 |
| 126 | SC inh. | F4 | Left | 82% | 78% | 20% | 0.41 |
| 21* | ACC-SC inh. | F4 | Left | 85% | 69% | 6% | 0.41 |
| 23* | ACC-SC inh. | F4 | Left | 92% | 77% | 5% | 0.35 |
| 103 | ACC-SC inh. | F4 | Left | 94% | 80% | 8% | 0.28 |
| 104 | ACC-SC inh. | F4 | Left | 98% | 93% | 14% | 0.36 |
| 110 | ACC-SC inh. | F4 | Left | 85% | 73% | 5% | 0.26 |
| 116* | ACC-SC inh. | F4 | Right | 75% | 65% | 3% | 0.35 |
| 38 | Imaging | F2,3 | Left | 83% | NA | 16% | 0.49 |
| 39 | Imaging | F2,3 | Left | 88% | NA | 19% | 0.35 |
| 41 | Imaging | F2,3 | Left | 84% | NA | 24% | 0.3 |
| 42 | Imaging | F2,3 | Left | 92% | NA | 20% | 0.37 |
| 43 | Imaging | F2,3 | Left | 76% | NA | 13% | 0.44 |
| 8 | VC inh. | F5-S1 | Left | 97% | 85% | 12% | 0.35 |
| 9 | VC inh. | F5-S1 | Left | 95% | 85% | 18% | 0.34 |
| 20 | VC inh. | F5-S1 | Left | 89% | 87% | 9% | 0.4 |
| 19 | VC inh. | F5-S1 | Left | 97% | 85% | 17% | 0.45 |
| 33 | VC inh. | F5-S1 | Left | 97% | 73% | 24% | 0.41 |
| 21* | ACC-VC<br>inh. | F5 | Right | 90% | 79% | 6% | 37% |
| 22 | ACC-VC<br>inh. | F5 | Right | 90% | 91% | 12% | 0.48 |
| 23* | ACC-VC<br>inh. | F5 | Right | 92% | 89% | 6% | 0.36 |
| 107 | ACC-VC<br>inh. | F5 | Left | 96% | 90% | 14% | 0.29 |
| 114 | ACC-VC<br>inh. | F5 | Right | 75% | 75% | 15% | 0.47 |
| 116* | ACC-VC<br>inh. | F5 | Right | 75% | 76% | 8% | 0.38 |
| 117* | ACC-SC<br>Rev. | F6 | Left | 81% | 71% | 6% | 0.35 |
| 118* | ACC-SC<br>Rev. | F6 | Left | 88% | 89% | 28% | 0.37 |
| 119* | ACC-SC<br>Rev. | F6 | Left | 71% | 69% | 12% | 0.42 |
| 127* | ACC-SC<br>Rev. | F6 | Right | 79% | 74% | 11% | 0.38 |
| 129* | ACC-SC<br>Rev. | F6 | Right | 76% | 68% | 18% | 0.38 |
| 153* | ACC-SC<br>Rev. | F6 | Right | 86% | 84% | 18% | 0.38 |
| 117* | ACC-VC<br>Rev. | F6 | Right | 81% | 80% | 22% | 0.38 |
| 118* | ACC-VC<br>Rev. | F6 | Right | 88% | 91% | 16% | 0.38 |
| 119* | ACC-VC<br>Rev. | F6 | Right | 71% | 72% | 6% | 0.36 |
| 127* | ACC-VC<br>Rev. | F6 | Left | 79% | 72% | 4% | 0.36 |
| 129* | ACC-VC<br>Rev. | F6 | Left | 76% | 71% | 16% | 0.41 |
| 153* | ACC-VC<br>Rev. | F6 | Left | 86% | 83% | 18% | 0.35 |

\* Mice contributed data to two experiments
<sup>a</sup> First figure referencing data from each mouse
<sup>b</sup> Accuracy on control trials from non-optogenetic sessions

### Activity of ACC-SC neurons during task performance

We used two-photon calcium imaging to determine how ACC activity relates to sensorimotor control during this task. We placed a chronic window over the ACC of Rbp4-Cre x Ai94D mice, which express the genetically encoded calcium sensor GCaMP6s in layer 5 excitatory neurons (Figure 2C). We analyzed responses of single neurons during four combinations of task cues and actions (contra-counterclockwise, contra-clockwise, ipsi-counterclockwise, and ipsi-clockwise). Individual neurons responded to multiple task variables, yet response amplitudes were modulated for specific cues and actions. For example, the neuron shown in Figure 2D selectively responded on contra cue trials but showed a higher response for clockwise vs. counterclockwise actions. The neuron in Figure 2E was active on both contra and ipsi cue trials as long as the animal selected a clockwise response. This pattern of activation also held at the population level; ACC activity was higher on clockwise than counterclockwise trials for both contra and ipsi cues (Figure 2F, G). We compared how ACC neurons respond to contra or ipsi cues without regard to the action selected by the animal. Overall, ACC neurons had higher activity on contra than ipsi cue trials (Figure 2H). We similarly compared responses on clockwise or counterclockwise trials regardless of which visual cue was presented. This showed that ACC neurons respond preferentially to clockwise actions (Figure 2H). These analyses suggest that, as a population, ACC neurons are preferentially activated by contra cues and clockwise actions during the task.

We performed pathway-specific imaging to determine whether and how ACC-SC neurons, identified by injecting a synthetic retrograde tracer into the SC (Figure 3A), respond to specific actions. Examination of single neuron and population responses showed higher activation of ACC-SC neurons on clockwise trials (Figure 3B-E). Next, we used a decoding approach to probe whether ACC-SC activity predicted selected actions (Figure 3F, G). Linear SVM classifiers trained with single-trial activity of ACC-SC neurons performed better than chance at predicting actions. Together, these analyses suggest that ACC neurons convey information about clockwise actions to the SC.

**Figure 3.**
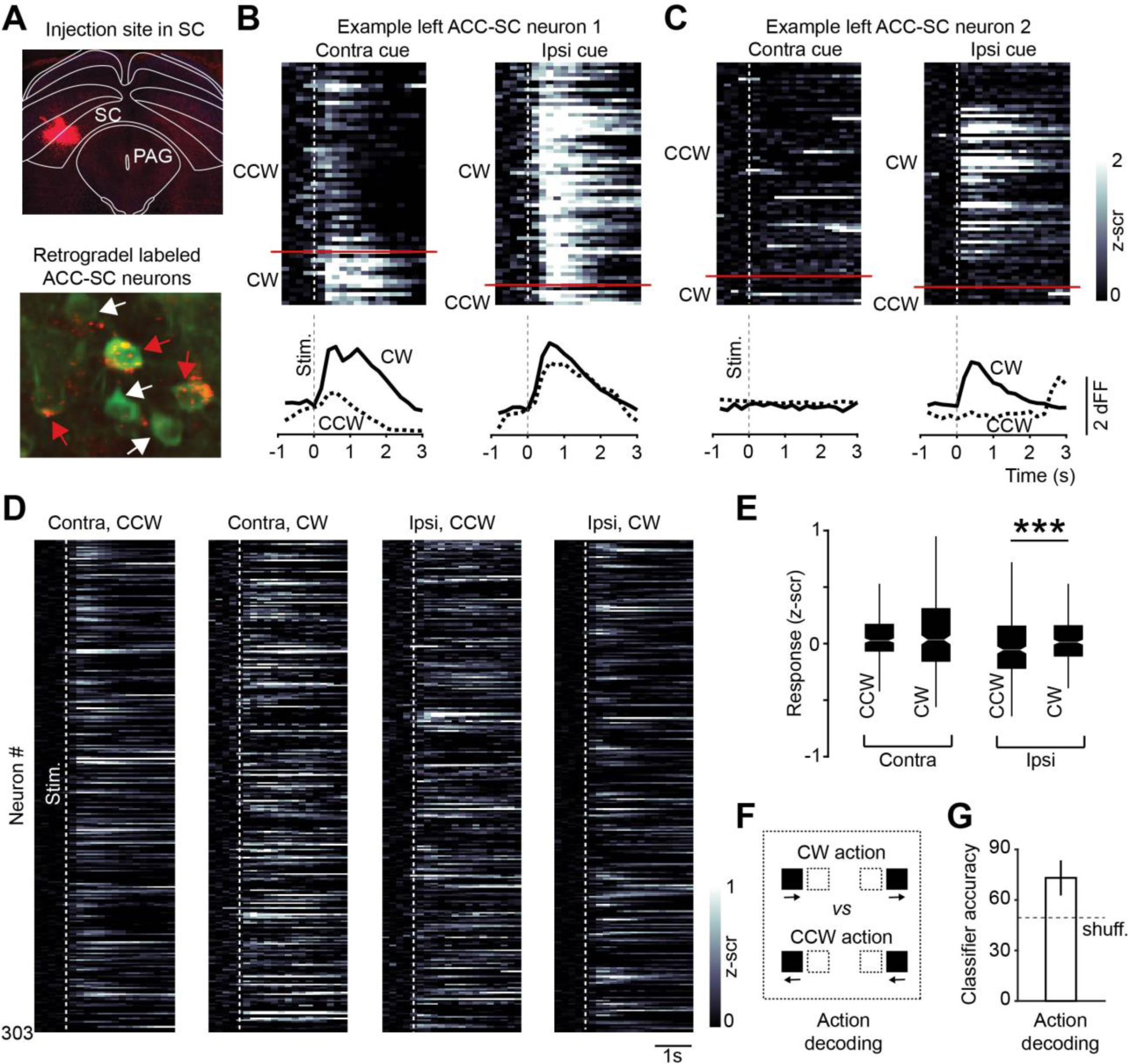
Task activity of ACC-SC neurons. (**A**) ACC-SC neurons were identified by injecting red retrobeads in SC_m_. Recordings were made from 303 ACC-SC neurons in five expert mice (8 behavioral sessions, 1-3 sessions per animal). (**B, C**) Task responses of two example ACC-SC neurons. Responses on contra and ipsi cue trials with counterclockwise or clockwise actions are shown. Rows show individual trials. Dotted lines show stimulus onset. (**D**) Session-averaged activity of individual ACC-SC neurons for the indicated trial type. Each row corresponds to the same neuron across the four plots. (**E**) Average responses on contra and ipsi cue trials with counterclockwise or clockwise responses (contra, CCW vs. CW: p = 0.300, z = -1.037; ipsi, CCW vs. CW: p = 0.001, z = -2.589). ***p < 0.005, Wilcoxon signed-rank test. (**F**) Combination of trials used for action decoding. SVM classifier was trained to distinguish responses on clockwise (right arrows) and counterclockwise trials (left arrows). Activity was combined as shown, without regard to the cue condition. (**G**) Classifier accuracy for action decoding. Dotted line shows chance model performance obtained by shuffling trial labels on the test set (see Methods). Error bar shows standard deviation of classifier accuracy over 1000 iterations of the model.

### The SC facilitates counterclockwise and the ACC-SC pathway clockwise actions

How does the clockwise action information observed in ACC-SC neurons contribute to task performance? We addressed this question by first examining the role of activity in the SC itself. We virally expressed the inhibitory opsin Jaws^52^ in the intermediate and deep layers of the left SC and delivered yellow light (593 nm) through an implanted optic fiber on a randomly selected subset of trials (Figure 4A). Illuminating the brain with light in the absence of an opsin did not produce significant changes in behavior, including in correct and incorrect responses on contra or ipsi cues (Supplementary Figure 2). To assess the effect of photoinhibition, we compared how mice responded to contra and ipsi cue trials under ‘laser’ and ‘no laser’ conditions (note that timeout trials, in which mice fail to give a complete response, are excluded from performance accuracy analysis and considered separately). Unilateral photoinhibition of the SC biased responses; inhibition increased incorrect responses on contra cue trials but decreased incorrect responses on ipsi cue trials (Figure 4B). In other words, SC inactivation decreased counterclockwise responses (Supplementary Figure 2A) and increased clockwise responses. These results are consistent with a recent neurophysiology study demonstrating that the activity of left SC neurons encodes counterclockwise actions in a similar task^53^. Reducing SC activity also increased the response time on contra cue trials and decreased the timeout rate on ipsi cue trials (Figure 4B). These results suggest that the left SC facilitates counterclockwise actions.

**Figure 4.**
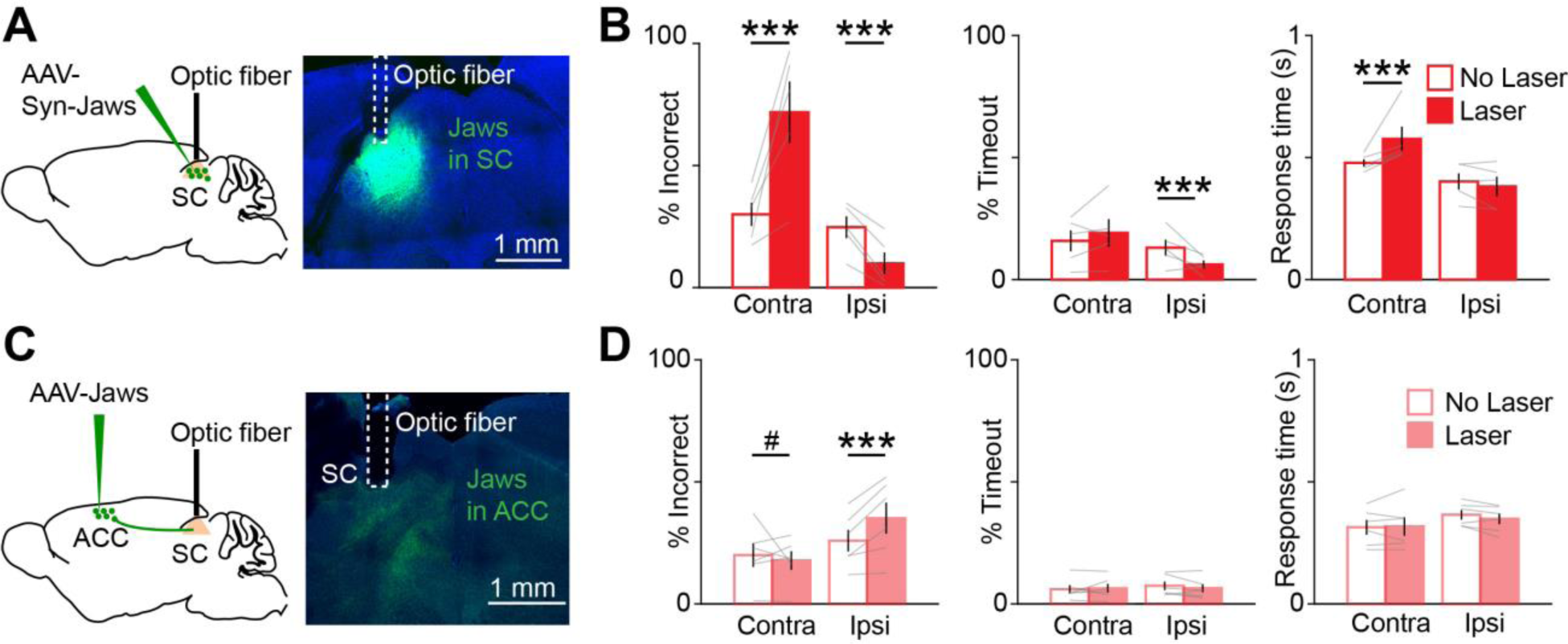
Effect of SC and ACC-SC inactivation on behavioral performance. (**A**) AAV5-Syn-Jaws was injected in the SC and a fiber optic was implanted above the injection site. (**B**) Behavioral performance for non-laser (unfilled) and laser (filled) conditions on contra and ipsi cue trials with SC inactivation (n = 5 mice). Incorrect performance (contra, p = 0.001; ipsi, p = 0.001), timeouts (i.e., trials with incomplete responses within the allotted time; contra, p = 0.077; ipsi, p = 0.001), and response time (contra, p = 0.001; ipsi, p = 0.025) are shown. ***p < 0.005; permutation test. (**C**) AAV5-CaMKII-Jaws was injected in the ACC and a fiber optic cannula was implanted in the SC. (**D**) Similar to B, except for inactivation of ACC outputs to SC (n = 6 mice). Incorrect performance (contra, p = 0.061; ipsi, p = 0.001), timeouts (contra, p = 0.961; ipsi, p = 0.561), and response time (contra, p = 0.198; ipsi, p = 0.092) are shown. #p = 0.061, ***p < 0.005; permutation test.

To determine if the left SC facilitates counterclockwise ball movements regardless of specific learned task contingencies, we tested a separate cohort of untrained mice as they spontaneously moved the ball. Inactivation of the left SC reduced counterclockwise and increased clockwise movements, which were like movements during the task (Supplementary Figure 3). The fact that SC inactivation decreases counterclockwise actions during spontaneous movement shows that this structure exerts an innate response bias and facilitates specific movement directions in head-fixed mice. These results, combined with the effects of SC inactivation during the visually guided task (Figure 4B), demonstrate that the left SC facilitates counterclockwise responses; by extension, the right SC facilitates clockwise responses.

How do direct ACC inputs to the SC modulate task performance? We addressed this question by virally expressing Jaws in the ACC and implanting an optic fiber cannula over the left SC to target ACC-SC inputs for inactivation (Figure 4C). ACC-SC inactivation increased incorrect responses on ipsi cue trials and there was a trend for decreased incorrect responses on contra cue trials (Figure 4D). That is, inactivation of left ACC-SC inputs increased counterclockwise actions and decreased clockwise actions. Hence, left ACC-SC inputs normally facilitate clockwise responses, consistent with the physiological responses of ACC-SC neuronal populations (Figure 3E-G). Given that the left SC promotes counterclockwise responses (Figure 4B; Supplementary Figure 3) and that each hemisphere of the ACC targets the SC on the same side (Figure 1H), these results suggest that the ACC-SC pathway promotes clockwise actions by reducing counterclockwise responses.

The SC facilitates counterclockwise responses (Figure 4B, Supplementary Figure 3); yet, unexpectedly, ACC inputs to the SC facilitate the opposite response (i.e., clockwise actions; Figure 4D). Since cortical projection neurons are predominantly glutamatergic, it is unclear how excitatory ACC-SC inputs implement a response bias opposite to the SC itself. We tested how the ACC influences activity in the SC; we photostimulated ChR2-expressing excitatory ACC neurons and used a 16-channel silicone probe to measure responses of SC neurons in the intermediate and deep layers of awake mice (Supplementary Figure 4A). Photostimulation modulated the activity of 27% of SC units (17/63 units from 3 mice) and led to a heterogeneous effect; while some SC units were excited, others were inhibited (Supplementary Figure 4B,C).

These optogenetic manipulations could exert their effect indirectly by recruiting polysynaptic inhibitory pathways through the basal ganglia^54^, excitatory-inhibitory commissural SC circuitry^55^, or other mechanisms. Alternatively, these results may reflect direct effects on SC neurons. We used anterograde trans-synaptic viral tracing^56^ to label SC neurons targeted by the ACC. We injected tdTomato reporter mice with an AAV1 virus expressing the Flpo recombinase in the ACC and performed immunohistochemistry against NeuN and GABA in slices containing the SC (Supplementary Figure 4D,E). Overall, 6.6 ± 1.3% of SC neurons expressed tdTomato (n = 5 mice). tdTomato labeling was observed in both GABA- and non GABA-containing neurons, suggesting that ACC inputs target both excitatory and inhibitory SC neurons. Yet, we found an overrepresentation of GABAergic neurons in our sample of tdTomato positive neurons, as compared to tdTomato negative neurons (Supplementary Figure 4F). Therefore, ACC neurons may recruit both excitatory and inhibitory neurons to modulate SC activity (Supplementary Figure 4G), potentially in a task-dependent manner.

### ACC outputs to the visual cortex facilitate performance on contra cue trials

Our results suggest that the ACC-SC pathway facilitates specific responses during the task. Sensory processing of behaviorally relevant stimuli is another key component of visual sensorimotor control. The ACC densely projects to the VC, modulates its activity, and facilitates performance on visually guided sensorimotor tasks^17, 18, 28, 29^. Hence, we determined the function of the ACC-VC pathway in this task. First, we performed retrograde tracing to determine the organization of ACC outputs to the VC and the SC. Injection of red and green fluorophore expressing rabies viruses in the SC and VC, respectively, showed that non-overlapping populations of ACC projection neurons target either structure, suggesting functional differentiation in ACC outputs (Figure 5A). To determine the contribution of the ACC-VC pathway, we first assessed the function of the VC in this task. Unilaterally inactivating the VC by delivering yellow light onto Jaws-expressing neurons increased incorrect responses on contra cue trials (Supplementary Figure 5). While this manipulation did not change the response time, there was an increase in the timeout rate on contra cue trials, consistent with a role for the VC in detection of contralateral visual cues^57^. These results suggest that the ACC may facilitate contra cue performance by modulating activity in VC. We directly tested this hypothesis by expressing Jaws in the ACC and photostimulating ACC output axons in the VC during the task (Figure 5B). Inactivating the ACC-VC pathway increased incorrect responses on contra cue trials (Figure 5C). Directly comparing the effect of inactivating the ACC-SC and ACC-VC pathways showed that ACC outputs to SC increased and outputs to VC decreased counterclockwise responses associated with the contra cue (Figure 5D). Thus, ACC-SC and ACC-VC pathways oppositely control behavioral performance in this task and may differentially modulate sensorimotor behavior.

**Figure 5.**
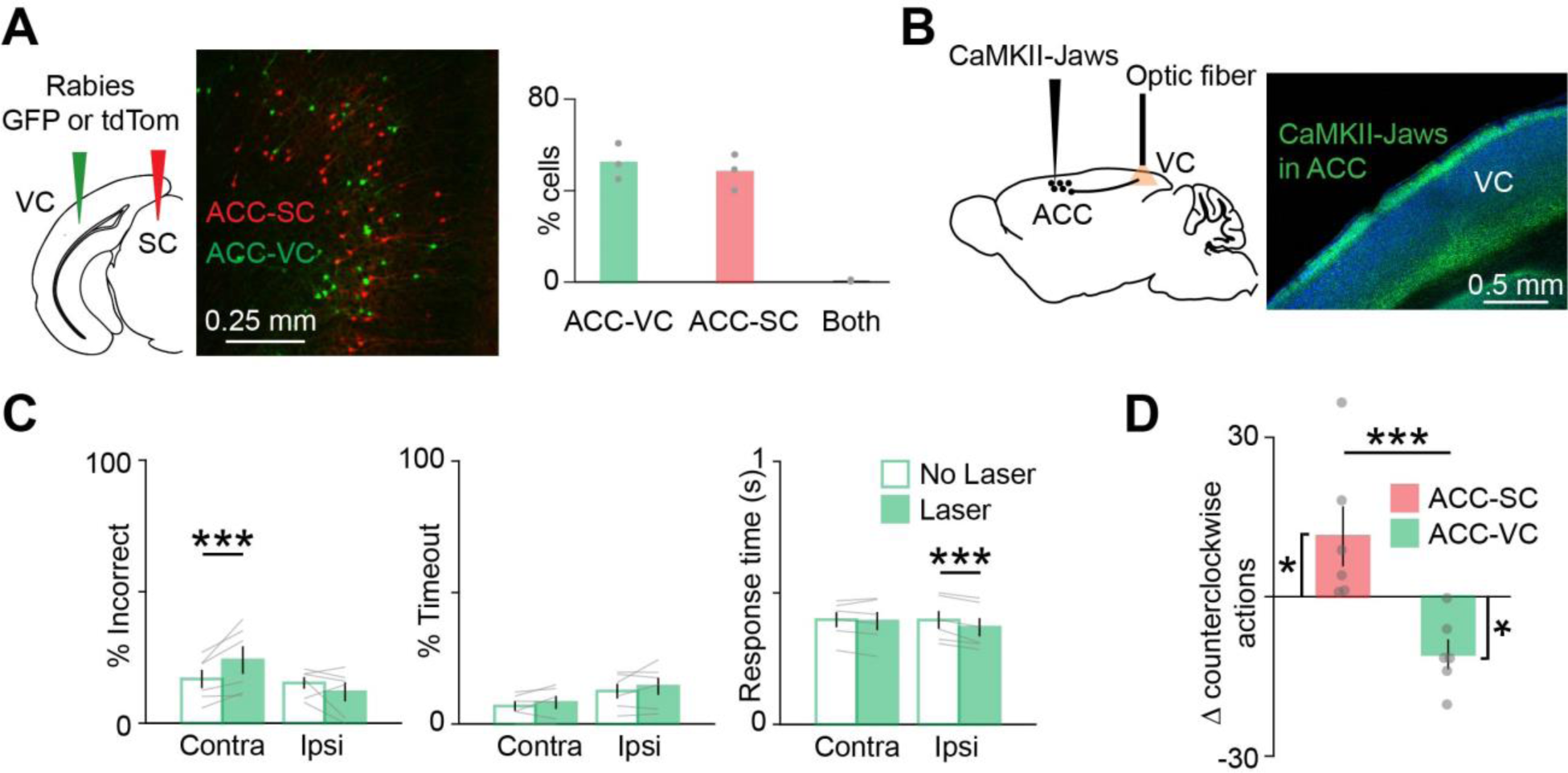
Effect of ACC-VC inactivation on behavioral performance. (**A**) *Left,* Dual-color retrograde tracing from VC and SC. Back-labeled ACC neurons projecting to SC_m_ (red) or VC (green). *Right,* Proportion of back-labeled neurons projecting to either or both areas (n = 3 mice). (**B**) AAV5-CaMKII-Jaws was injected in the ACC and axons in VC were targeted for inactivation. (**C**) Behavioral performance for non-laser (unfilled) and laser (filled) conditions on contra and ipsi cue trials with inactivation of ACC outputs to the VC (n = 6 mice). Incorrect performance (contra, p = 0.002; ipsi, p = 0.153), timeouts (contra, p = 0.204; ipsi, p = 0.312) and response time (contra, p = 0.656; ipsi, p = 0.006) are shown. (**D**) Comparison of normalized laser-induced change in counterclockwise actions with inactivation of ACC outputs to SC and VC (ACC-SC vs. zero, n = 6 mice, p = 0.046, z = -1.992; ACC-VC vs. zero, n = 6 mice, p = 0.028, z = 2.201; ACC-SC vs. ACC-VC, p = 0.003, z = 2.802). Significance testing with permutation test in C; in D, comparison against zero with two-tailed Wilcoxon signed-rank test and comparison between ACC-SC and ACC-VC with one-tailed Wilcoxon rank-sum test. Error bars are standard error of the mean.

### ACC-SC and ACC-VC pathways differentially modulate sensory processing and motor responses

How do ACC-SC and ACC-VC pathways control task performance? Optogenetic inactivation of these pathways can directly modulate specific responses or disrupt processing of visual stimuli that cue responses. We dissociated these possibilities by training a new cohort of mice on a reversed sensorimotor contingency, such that mice were required to move the presented visual cue outward to the side of the screen instead of the center (Figure 6A). In this task, clockwise responses previously associated with an ipsi stimulus were cued by the contra stimulus (and vice versa). Inactivation of the ACC-SC pathway caused a reversal in deficit and increased incorrect responses on contra cue trials (Figure 6B), as opposed to the deficit observed on ipsi cue trials in the previous task contingency (Figure 4D). Thus, the ACC-SC pathway facilitates clockwise responses in either task contingency. In contrast, inactivating the ACC-VC pathway increased incorrect responses on contra cue trials (Figure 6C), similar to that observed in the previous task (Figure 5C). Hence, the ACC-VC pathway is important for sensory processing of contra stimuli in both tasks. We did not observe significant changes in timeouts or the reaction time with either manipulation in the outward task (Figure 6B, C).

**Figure 6.**
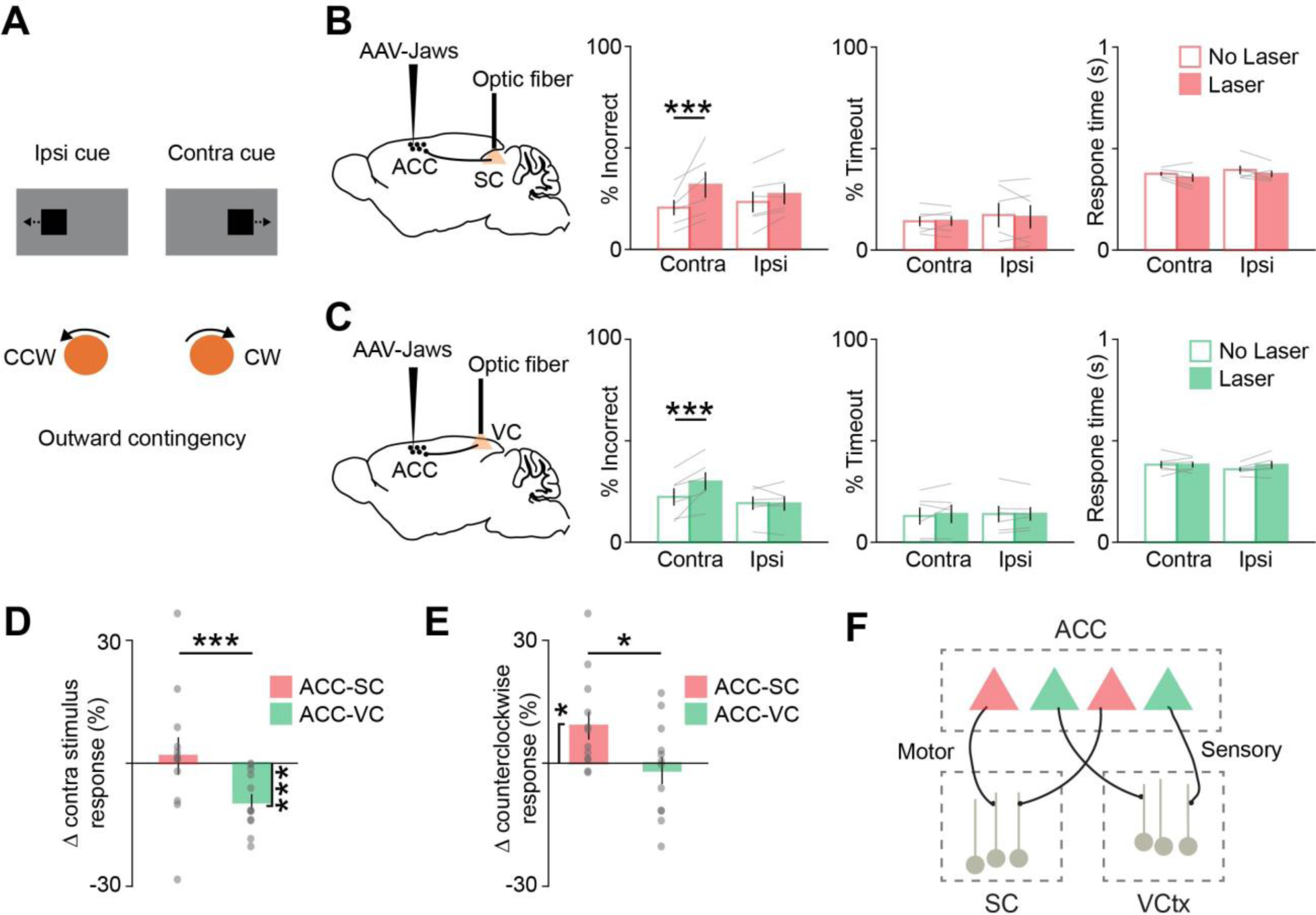
ACC outputs to SC and VC differentially modulate motor and sensory processing, respectively. (**A**) Mice were trained on the outward sensorimotor contingency and were required to move ipsi and contra cues away from the center by rotating the trackball counterclockwise and clockwise, respectively. Note that in this task, a counterclockwise action is the correct response for ipsi cues but an incorrect response for contra cues. (**B**) Behavioral performance for non-laser (unfilled) and laser (filled) conditions on contra and ipsi cue trials with inactivation of ACC outputs to the SC in the outward task (n = 6 mice). Incorrect performance (contra, p = 0.001; ipsi, p = 0.127), timeouts (contra, p = 0.924; ipsi, p = 0.927), and response time (contra, p = 0.033; ipsi, p = 0.138) are shown. ***p < 0.005; permutation test. (**C**) Similar to B, except for inactivation of ACC outputs to the VC (n = 6 mice). Incorrect performance (contra, p = 0.002; ipsi, p = 0.786), timeouts (contra, p = 0.499; ipsi, p = 0.885), and response time (contra, p = 0.869; ipsi, p = 0.042) are shown. ***p < 0.005; permutation test. (**D**) Data from both tasks was combined. Effect of ACC-SC (n = 12 mice) and ACC-VC inactivation (n = 12 mice) on laser-induced change in contra stimulus responses (ACC-SC vs. zero, p = 0.638, z = -0.471; ACC-VC vs. zero, p = 0.002, z = 3.059; ACC-SC vs. ACC-VC, p = 0.004, z = 2.685). ***p < 0.005; comparison against zero with two-tailed Wilcoxon signed-rank test, and comparison between ACC-SC and ACC-VC with one-tailed Wilcoxon rank-sum test. (**E**) Similar to D, except showing laser-induced change in counterclockwise responses (ACC-SC vs. zero, p = 0.019, z = -2.353; ACC-VC vs. zero, p = 0.695, z = 0.392; ACC-SC vs. ACC-VC, p = 0.03, z = 1.876). *p < 0.05; comparison against zero with two-tailed Wilcoxon signed-rank test, and comparison between ACC-SC and ACC-VC with one-tailed Wilcoxon rank-sum test. (**F**) Model summarizing the sensory and motor roles of ACC projections to the VC and SC, respectively.

We further determined how deficits observed with ACC-SC and ACC-VC pathways map onto responses to specific visual cues or the direction of the response itself. We combined data across the two tasks and computed a contra stimulus response index to compare how inactivation of ACC-SC and ACC-VC pathways affects responses associated with the contralateral visual cue. Inactivation of the ACC-VC pathway significantly decreased responses cued by contra stimuli, whereas inactivation of the ACC-SC pathway had no consistent effect (Figure 6D). Next, we computed a counterclockwise response index to determine how optogenetic inactivation modulates counterclockwise responses regardless of which stimulus cues them. While ACC-SC inactivation increased counterclockwise responses, we did not observe a significant change with ACC-VC inactivation (Figure 6E). Thus, the ACC-SC pathway mediates specific motor responses, whereas the ACC-VC pathway facilitates sensory processing in these tasks (Figure 6F).

## Discussion

We demonstrate that the ACC modulates visual sensorimotor behaviors by using anatomically distinct but functionally complementary populations of projection neurons to facilitate sensory processing (ACC-VC) and specific motor responses (ACC-SC; Figure 6D-F). We show that VC inputs bring contralateral stimulus information and callosal inputs bring ipsilateral stimulus information to the caudal ACC (Figure 1A-G; Supplementary Figure 1). In turn, a subset of ACC neurons projects to the SC (Figure 1H). Using a two-choice visual sensorimotor task, we show that the activity of ACC-SC neurons predicts actions and responds to clockwise movements (Figure 3E,G). SC inactivation decreases counterclockwise responses during the inward task and spontaneous movements (Figure 4A,B; Supplementary Figure 3), consistent with a role for this area in response selection^9, 37^. Surprisingly, inactivation of the ACC-SC pathway has the opposite effect and disrupts performance by decreasing clockwise responses (Figure 4D). Importantly, by training mice on a reversed sensorimotor contingency (outward task), we demonstrate that ACC-SC inactivation consistently decreases clockwise responses regardless of the specific cue-response mapping (Figure 6B,E). Hence, the ACC-SC pathway primarily contributes to motor responses. In contrast, ACC-VC inactivation decreases performance on contra cue trials in either task (Figure 5C, 6C), suggesting that it facilitates contra cue processing regardless of the associated response.

The SC is an integrative node within the broader midbrain selection network, serving as a key arbiter for response selection across multiple sensory and motor modalities in many species^9, 32–36, 42, 58^. In our task, mice make responses by rotating a ball with their forepaws. Our finding that SC activity crucially contributes to such responses provides further evidence that the SC plays a general role in the selection of directional responses beyond eye and whole-body movements. We find that unilateral SC inactivation does not increase inaction, but rather changes the likelihood of selecting specific responses (Figure 4B). In other left-right response tasks, neurophysiological recordings show a subset of SC neurons respond maximally for contralateral responses (i.e., movements directed opposite to the side of hemisphere under study) and are often silent or inhibited for ipsilateral responses^9, 53, 59^. Similarly, causal activity manipulations produce response biases consistent with these activity measurements^41, 42, 59^. These results, combined with our own, suggest a ‘winner-take-all’ interhemispheric competition model for selection in midbrain circuits, wherein the final response is determined by the SC hemisphere with the higher level of activity^9, 60^. Although the competition process underlying response selection likely involves an interplay between multiple systems, including intrinsic and commissural SC circuitry^58, 61, 62^ and inputs from the basal ganglia^54^, our results identify the important role of direct ACC projections to the SC.

We found that SC and ACC-SC inactivation oppositely modulates behavioral performance in the inward task contingency (Figure 4B, 4D). Several mechanisms could mediate this effect. In untrained mice, photostimulation of ACC neurons both excites and inhibits a subset of SC units (Supplementary Figure 4A-C). Moreover, ACC outputs directly target inhibitory as well as excitatory SC neurons (Supplementary Figure 4D-F). Hence, one possibility is that ACC projections recruit local inhibition in the left SC on ipsi cue trials (Supplementary Figure 4G), tipping the balance in favor of activity in the right SC and increasing the probability of clockwise responses. Alternatively, this effect may be mediated via commissural excitatory SC neurons^61, 62^ that are targeted by ACC neurons. In this case, during an ipsi cue trial, activation of left ACC-SC neurons would indirectly increase activity in the right SC (in addition to the ipsi cue directly activating the SC via inputs from the overlying sensory layer and other structures), thereby leading to a clockwise response. A better understanding of the downstream anatomical targets of SC neurons that receive ACC inputs will help clarify these issues. Regardless of exact mechanisms, we clearly demonstrate opposing influence of ACC-SC and SC computations on response selection.

A subset of ACC-SC neurons was recently shown to provide collaterals to the lateral posterior (LP) thalamus and modulate cortical sensory processing in the VC^30^. This raises the possibility that ACC-SC neurons are anatomically and functionally heterogeneous. While we clearly demonstrate that direct ACC-SC outputs are critical for facilitating specific motor responses and contribute minimally to sensory processing, it is possible that the sensory function of this pathway was not engaged by the relatively simple visual stimuli used in this task. In contrast to the inward task, we found that the ACC-SC pathway is important for responses on contra cue trials in the outward task contingency (Figure 6B). Hence, task responses of ACC-SC neurons are likely shaped by specific sensorimotor contingencies.

Our results also support the hypothesis that the PFC exerts executive control over task responses by modulating activity in downstream motor structures. In the inward task, mice move their forelimbs in a direction that would orient them to the visual cue if they were freely moving, potentially similar to a ‘pro’ response in eye movement or whole-body orienting tasks^43, 63^. In the outward task contingency, mice make movements that orient them away from the visual cue, possibly similar to an ‘anti’ response. The PFC is thought to play a crucial role in facilitating ‘anti’ performance^63^. Our projection-specific inactivation of the ACC-SC pathway in the inward and outward task contingencies (Figures 4D; 6B, E) are consistent with a role for the PFC in facilitating ‘anti’ performance.

Although ACC outputs to the SC coordinate specific responses, we found that the ACC uses an anatomically non-overlapping but functionally complementary population of projection neurons to facilitate sensory processing through the visual cortex (Figures 5C; 6C,D). An important issue is whether ACC outputs to the VC and SC control behavioral performance by modulating sensory or motor components of the task. Previous studies have addressed this issue using delay tasks with stimulus presentation and motor response epochs separated by an intervening delay^2, 5, 64, 65^. However, in such tasks, the cue-response mapping is known to the subject before stimulus presentation and movement planning can start with stimulus presentation^66^, making it difficult to disambiguate whether temporally restricted inactivation affects behavior by disrupting sensory or motor processing. We instead addressed this issue by training a set of mice on the outward sensorimotor contingency and comparing the effect of projection-specific inactivation across task conditions. This showed that the ACC-VC pathway is important for responses associated with contralateral visual cues (Figure 6C, D), suggesting it predominantly contributes to sensory processing. This is consistent with previous work demonstrating that the ACC-VC pathway modulates stimulus encoding by VC neurons in passively-viewing mice and facilitates stimulus discrimination^17^. While the exact mechanisms mediating task performance through the visual cortex are unclear, the ACC-VC projection may control VC outputs to multiple pathways that ultimately converge in the SC and other parts of the midbrain selection network^54,67,68^.

Overall, our results highlight the importance of projection-specific manipulations in multiple task contingencies for dissecting circuit mechanisms underlying sensorimotor behaviors. By dissociating the contribution of individual outputs, our findings suggest a general organizing principle for PFC circuits wherein complementary behavioral functions are fulfilled by anatomically distinct output pathways, thereby enabling independent control over specific functions depending on task demands.

---

**Supplementary Figure 1.**
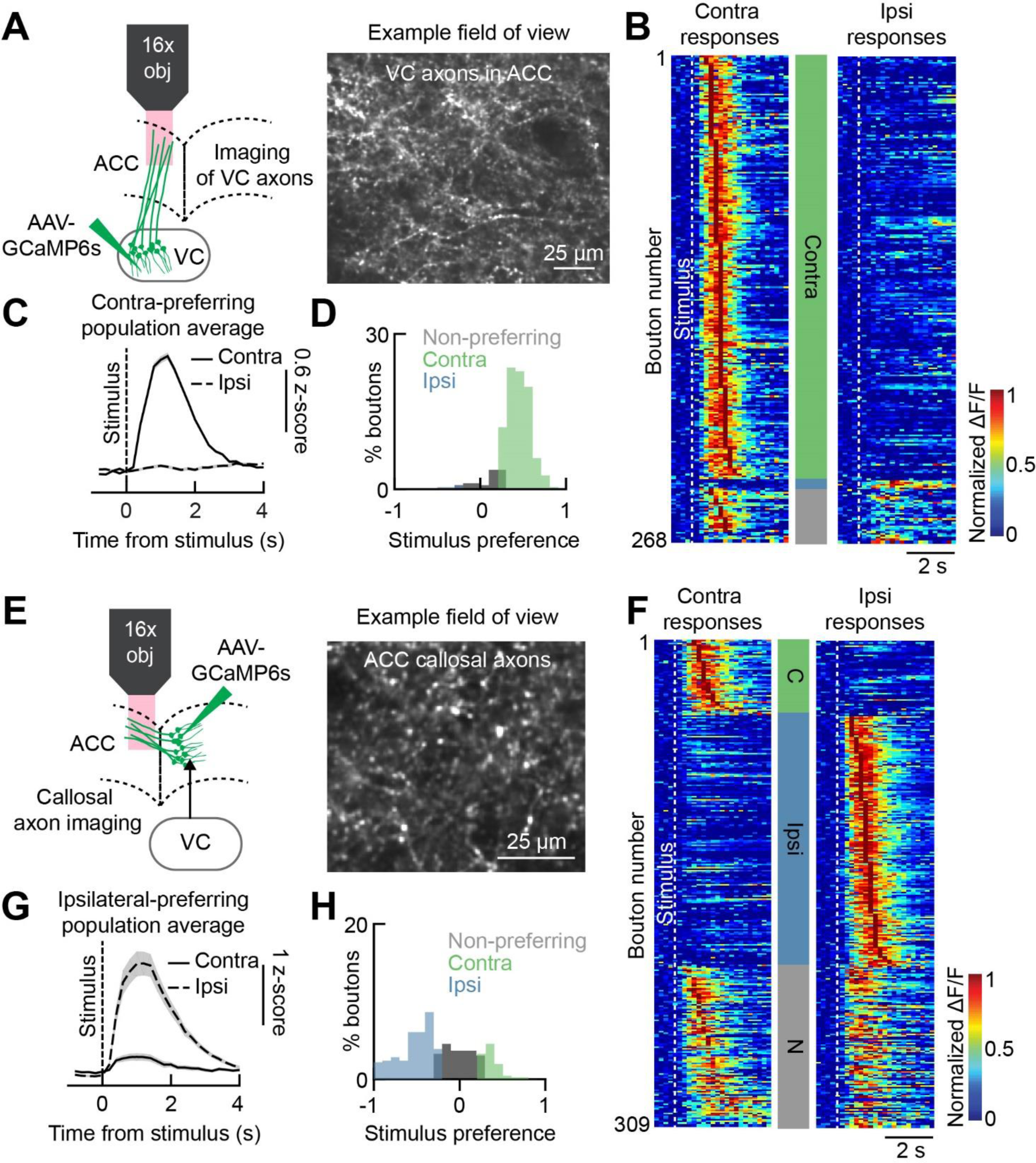
Visually-evoked responses of VC and callosal inputs to the ACC. (**A**) Two-photon calcium imaging of GCaMP6s-expressing VC axons in the ACC. (**B**) Session-averaged responses of individual visually-responsive boutons to stimuli presented in hemifield contra- or ipsilateral to the recording site (black square, ∼20° size, 1s). Responses are grouped by their stimulus preference (contralateral, green; ipsilateral, blue, non-preferring, gray) and sorted within each group by their peak response time. (**C**) Population-averaged response of contra-preferring boutons to contralateral (solid line) and ipsilateral (dashed line) stimuli. Shading shows the standard error of the mean. (**D**) AUROC analysis was used to calculate a stimulus preference score for each visually-responsive bouton. This score ranges from -1 (ipsi selective) to 1 (contra selective). Distribution of scores for 268 visually-responsive VC boutons in ACC from 4 mice is shown. (**E-H**) Same as **A-D**, except for callosal boutons (n = 309 boutons from 5 mice).

**Supplementary Figure 2.**
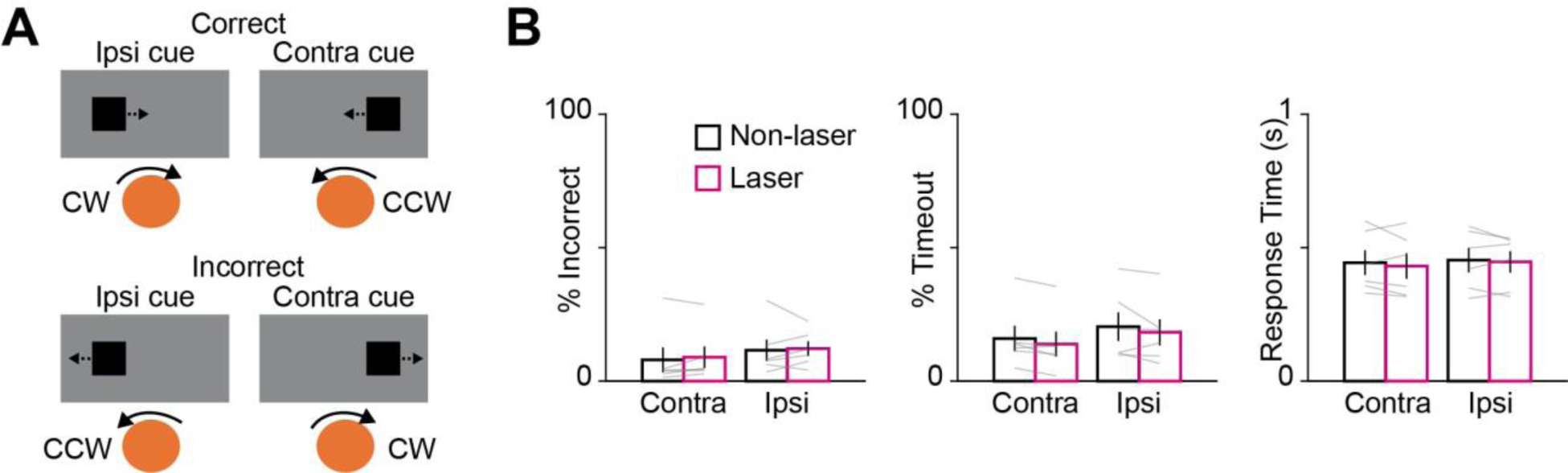
Effect of light delivery on behavioral performance. (**A**) Schematic depicting correct and incorrect responses on the inward task. (**B**) Behavioral performance with light delivery (20mW) in the absence of opsins (n = 6 mice). Behavioral performance for non-laser and laser conditions on contra cue and ipsi cue trials. Incorrect performance (contra, p = 0.201; ipsi, p = 0.842), timeouts (contra, p = 0.998; ipsi, p = 0.078), and response time (contra, p = 0.527; ipsi, p = 0.647) are shown. Significance testing with permutation test.

**Supplementary Figure 3.**
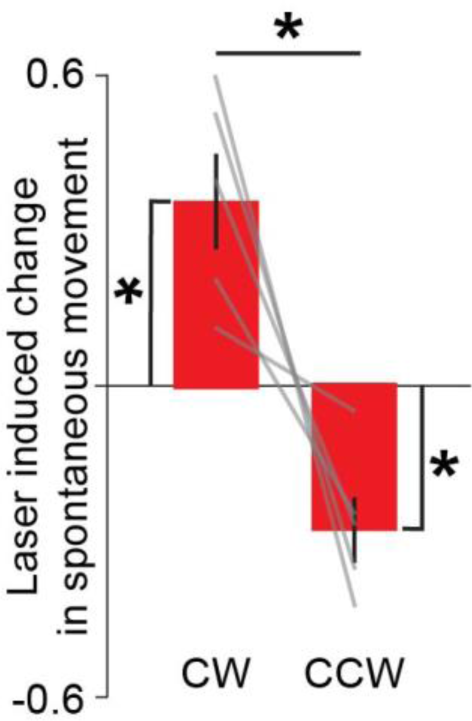
SC inactivation during spontaneous movement. AAV5-Syn-Jaws was injected in the SC and a fiber optic was implanted over the injection site. Laser induced changes in clockwise and counterclockwise actions during spontaneous movements are shown (CW vs. zero: p = 0.018; CCW vs. zero: 0.012; CW vs. CCW: 0.014; n = 5 mice). *p < 0.05; one-sample t-test against zero, and two sample t-test for comparison between actions. Error bars are standard error of the mean.

**Supplementary Figure 4.**
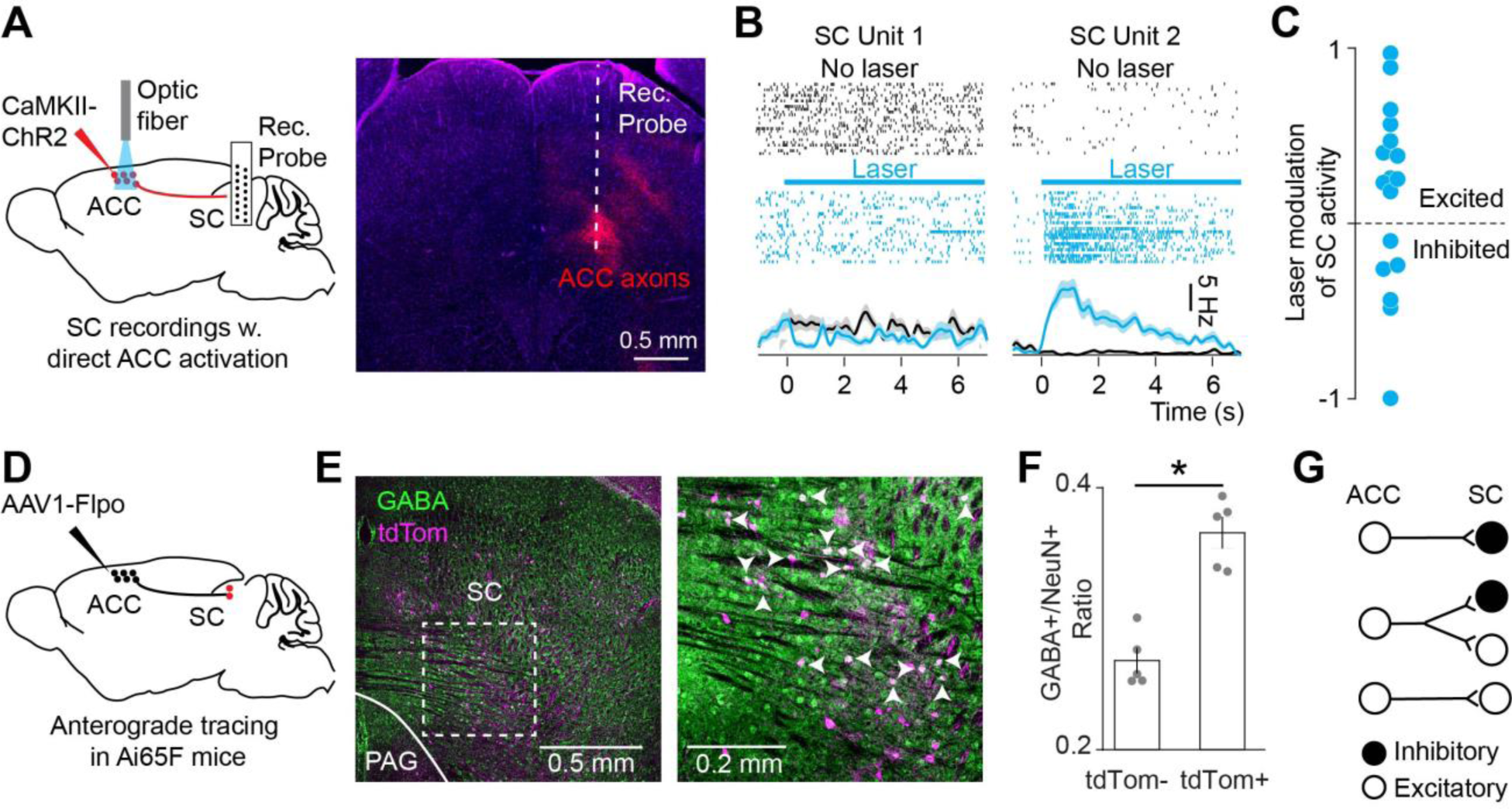
Effect of ACC activation on SC activity. (**A**) AAV virus encoding CaMKII-ChR2-mCherry was injected unilaterally in the ACC and an optic fiber was implanted above the injection site. Extracellular recording probe was inserted in the SC on the same side as the ACC optic fiber. (**B**) Activity of two example SC units with and without photostimulation of the ACC. (**C**) Laser-modulation indices for SC units significantly modulated with ACC photoactivation. (**D**) SC neurons receiving inputs from the ACC were labeled by injecting AAV1-Flpo virus in the ACC of tdTomato reporter mice. (**E**) *Left,* SC neurons labeled with tdTomato (magenta). Imunnohistochemistry against GABA (green) shows inhibitory neurons. *Right,* higher magnification image of the area bounded by the dotted white square in left panel. Arrowheads mark neurons double-labeled with tdTomato and GABA. (**F**) Proportion of tdTomato unlabeled (tdTom-) and labeled (tdTom+) neurons (i.e., NeuN positive cells) that are co-labeled with GABA (*p = 0.0431, z = -2.023). Significance testing with two-tailed Wilcoxon signed-rank test. Error bars are standard error of the mean. (**G**) Proposed circuit schematic for ACC connectivity with excitatory and inhibitory SC neurons.

**Supplementary Figure 5.**
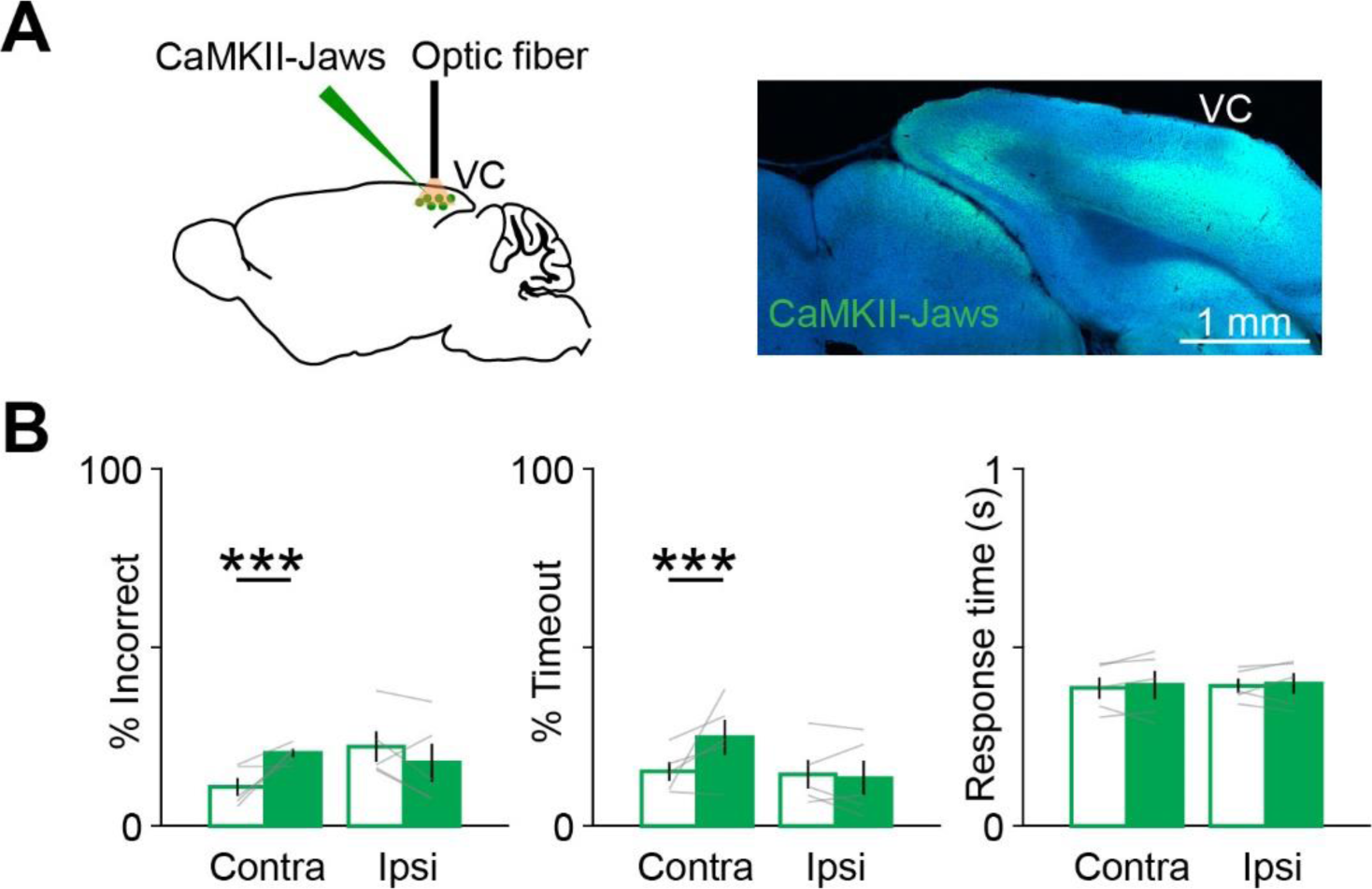
Effect of VC inactivation on task performance. (**A**) AAV5-CaMKII-Jaws was injected in the VC, which was inactivated via yellow light through a chronic window. (**B**) Behavioral performance for non-laser (unfilled) and laser (filled) conditions on contra and ipsi cue trials with inactivation of the VC (n = 5 mice). Incorrect performance (contra, p = 0.001; ipsi, p = 0.117), timeouts (contra, p = 0.001; ipsi, p = 0.496), and response time (contra, p = 0.232; ipsi, p = 0.966) are shown. Significance testing with permutation test. Significance evaluated at the Bonferroni-adjusted α-value of 0.025. ***p < 0.005.

## Acknowledgments

We thank Michael Halassa for thoughtful discussions and Vincent Pham for assistance with histology experiments. This work was supported by grants from National Eye Institute F32 EY024857 to R.H., National Institute of Mental Health K99 MH112855 to R.H., National Eye Institute F32 EY028028 to G.O.S., Natural Sciences and Engineering Research Council of Canada PDF 487824-2016 to V.B.P., National Institute of Mental Health U01 MH106018 and U01 MH109129 to I.R.W., National Eye Institute R01 EY007023 to M.S., National Institute of Neurological Disease and Stroke U01 NS090473 to M.S., National Science Foundation EF1451125 to M.S., and Simons Foundation Autism Research Initiative to M.S.

## Author contributions

R.H. and M.S. conceived the project and developed the concepts presented. R.H. designed the experiments and analysis approach with contribution from M.S. R.H. and G.P. designed and implemented the behavioral task. R.H. performed and analyzed the anatomical, behavioral, optogenetic, and calcium imaging experiments with contributions from G.O.S., K.G.C., L.G., A.S., and E.A. V.B.P. performed and analyzed the extracellular recording experiments with input from R.H. I.C.W. provided unpublished reagents. R.H. and M.S. wrote the manuscript with input from all authors.

## Declaration of interests

The authors declare no competing interests.

## Methods

### Animals

All experimental procedures performed on mice were approved by the Massachusetts Institute of Technology Animal Care and Use Committee. Mice were housed on a 12 hour light/dark cycle. Animals were group-housed before surgery and singly-housed afterwards. Adult mice (> 2 months) of either sex were used for these studies. In addition to wild type mice, the following transgenic lines were used: Rbp4-Cre (MMMRC stock # 031125), Ai94(TITL-GCaMP6s)-D (Jackson Laboratory stock # 024104), and Ai65(RCFL-tdT)-D (Jackson Laboratory stock # 021875).

### Surgical procedures

Surgeries were performed under isoflurane anesthesia (3-4% induction, 1-2.5% maintenance). Animals were given analgesia (slow release buprenex, 0.1mg/kg and Meloxicam 0.1mg/kg) before surgery and their recovery was monitored daily for 72 hours. Once anesthetized, animals were fixed in a stereotaxic frame. The scalp was sterilized with betadine and ethanol. For anatomical tracing experiments, we made a midline incision in the scalp using a scalpel blade. Depending on the experiment, rabies or AAV viruses were injected in the visual cortex (AP: -3.5mm, ML: 2.5mm, DV: 0.5 mm), caudal ACC (AP: 0.3mm, ML: 0.5mm, DV: 0.5 mm), rostral ACC (AP: 1.6mm, ML: 0.3mm, DV: 1.3 mm), or the superior colliculus (at AP: - 3.6mm, ML: 1mm, and DV: 1.5mm) using a microinjector (Stoelting). After virus injection, the scalp was reclosed with sutures and skin adhesive (Vetbond).

The following surgical procedures were performed for optogenetics or imaging experiments. For experiments requiring the use of dental acrylic (fiber optic cannulae and chronic imaging windows), we removed a portion of the scalp using spring scissors, scraped away the periosteum membrane overlying the skull, and used a dental drill to abrade the skull to improve adhesion. For light control experiments, two mice expressed axonal GCaMP6s and four mice were wildtype (Supplementary Figure 2B). For photoinhibition of the superior colliculus (SC) during the inward task or spontaneous movements, we injected AAV5-hSyn.Jaws-KGC-GFP-ER2 (100nL) in the intermediate/motor layer (AP: -3.6mm, ML: 1mm, DV: 1.5mm) and implanted a fiber optic cannula (300µm/0.39NA core, CFM13L02, Thorlab) 0.2mm dorsal to the injection site. Cannulae were secured on the skull using layers of Metabond and were protected with a dust cap until used for experiments. For modulating ACC outputs to the superior colliculus, we injected AAV5-CaMKII-Jaws-KGC-GFP-ER2 (University of North Carolina vector core) in the ACC and implanted a fiber optic cannula (300µm/0.39NA core or 400µm/0.5 NA core, CFMXD02, Thorlabs) over the intermediate layer of the superior colliculus (AP: -3.6mm, ML: 1mm, DV: 1.3mm). For optogenetic inactivation of the visual cortex (VC), we drilled a 3mm craniotomy and made 8-12 injections (100nL each) of an AAV5.CaMKII.Jaws-KGC-GFP-ER2 (University of North Carolina vector core) virus 0.5mm below the surface in a grid pattern. Injections were centered on the left primary visual cortex (centered at AP: -3.5mm, ML: 2.5mm; range AP: 4 to 3mm, ML: 2.2 to 2.8mm). For inactivation of ACC axons in VC, we injected AAV5-CaMKII-Jaws-KGC-GFP-ER2 in the ACC and implanted a 3mm chronic window or a fiber optic cannula (400µm/0.5 NA core) over the VC (centered at AP: -3.5mm, ML: 2.5mm, AP = 0, on pia). In a subset of experiments on the inward task, AAV5-CaMKII-Jaws-KGC-GFP-ER2 was injected bilaterally in the ACC and the VC or SC was targeted for axonal inactivation on either side of the brain. For projection-specific optogenetic inactivations during the outward contingency task, we bilaterally injected AAV5-CaMKII-Jaws-KGC-GFP-ER2 in the ACC and implanted fiber optic cannulas over the SC and VC on either side of the brain (see Table 1 for details on which brain hemisphere was targeted for each experiment).

For all imaging experiments in the ACC, we drilled a 3mm craniotomy over the midline (centered at AP: 0.5mm and ML: 0mm) and implanted a chronic imaging window assembled from two 3mm coverslips glued to a 5mm coverslip using a UV curable adhesive (Norland 61; double-chronic window). Two 3mm coverslips were required to minimize movement artifacts for GCaMP recordings during task performance. To label superior colliculus-projecting ACC neurons (ACC-SC), we injected 100nL of red retrobeads (Lumaflour) in the intermediate/motor layer of the SC (at AP: -3.6mm, ML: 1mm, DV: 1.7mm). For axonal imaging experiments, 200nL of AAV1-Syn-GCaMP6-WPRE-SV40 virus was injected unilaterally in the visual cortex (AP: -3.5mm, ML: 2.5 mm, DV: 0.5 mm) or the opposite ACC (AP: 0.5mm, ML: 0.5 mm, DV: 0.5 mm).

For all experiments, after implantation of chronic windows or optic fiber cannulas, the skull was attached to a stainless-steel custom-designed head plate (eMachines.com) using Metabond. Animals were allowed to recover for at least five days before commencing water restriction for behavioral experiments.

### Virus-mediated anatomical tracing

Standard histological techniques were used for analysis of retrograde and anterograde transsynaptic tracing experiments and for post-hoc verification of implantation/injection sites. Mice were deeply anesthetized with isofluorane and transcardially perfused with a 4% paraformaldehyde (PFA) solution prepared in phosphate buffered saline (PBS).

Extracted brains were postfixed in 4% PFA overnight at 4°C, then kept in PBS until sectioning. Fixed brain tissue was sectioned using a microtome (Leica VT-1000) into coronal slices (thickness of 50-100 µm, depending on the experiment). Slices were stained with DAPI and mounted on glass microslides using Vectashield hardset mounting media. Mounted sections were stored at 4°C until they were imaged using a laser scanning confocal microscope (Leica SP8).

Rabies viral vectors were made as described^69^. Briefly, HEK 293T cells (ATCC CRL-11268) were transfected with expression vectors for the ribozyme-flanked viral genome (cSPBN-4GFP (Addgene 52487) or pRVΔG-4tdTomato (Addgene 52500)), rabies viral genes (pCAG-B19N (Addgene 59924), pCAG-B19P (Addgene 59925), pCAG-B19G (Addgene 59921), and pCAG-B19L (Addgene 59922)), and the T7 polymerase (pCAG-T7Pol (Addgene 59926)). Supernatants were collected from 4 to 7 days after transfection, filtered, and pooled, passaged 3-4 times on BHK-B19G2 cells ^70^ at a multiplicity of infection of 2-5, then passaged on BHK-EnvA2 cells at a multiplicity of infection of 2. Purification, concentration, and titering were done as described.

To make AAV1-CAG-Flpo virus, the Flpo gene ^71^ was cloned into pAAV-CAG-FLEX-EGFP (Addgene 59331) to make pAAV-CAG-Flpo. Serotype 1 AAV was produced by triple transfection of HEK 293T cells with pAAV-syn-Flpo, pAAV-RC1, and pHelper (Cellbiolabs VPK-401) (per 15 cm plate, 15.5 ug, 21.0 ug, and 33.4 ug respectively) using Xfect (Clontech 631318) transfection reagent. Supernatants and cells were harvested at 72 hours posttransfection and viruses purified and concentrated by iodixanol gradient centrifugation^72^.

For quantification of dual-rabies retrograde tracing experiments from caudal and rostral ACC, we sliced the brain into 100 µm sections and imaged GFP and/or tdTomato fluorescence from every other slice using a 10x/0.4NA objective (Leica). Anatomical landmarks, such as position/size of ventricles, corpus callosum, striatum, and hippocampus, were used to manually align slices to a standard mouse brain atlas^46^. Back-labeled cells were counted using the cell counter plugin in ImageJ (NIH). For the results presented in Figure 1C, we counted the number of retrogradely neurons present in the different divisions of the visual cortex and normalized by the total number of GFP- and tdTomato-expressing back-labeled cells found in the posterior cortex (0 to -4.0 mm posterior, relative to Bregma). For quantifying the distribution of cells projecting to the intermediate and deep layers of the SC (Figure 1I), we counted the number of cells found in the ACC as a function of distance from Bregma in 0.5mm bins. Since the sensory and intermediate/motor layers of the superior colliculus are <0.5mm apart, it is challenging to restrict virus injection to the motor layer. However, the frontal cortex predominantly projects only to the motor layer; hence, to minimize contamination from virus spillover into the sensory layer of the superior colliculus, we normalized the number of neurons in the ACC by the total number of back-labeled neurons found in the frontal cortex (0 to +3mm anterior, relative to Bregma).

We injected rabies viruses encoding GFP or tdTomato into the visual cortex or the superior colliculus to identify ACC neurons projecting to these structures. We counted the total number of back-labeled neurons in the ACC (AP range 0 to 1mm) every 200 µm in each animal and quantified the proportion of neurons that were labeled with GFP, tdTomato, or both out of all labeled cells.

We performed anterograde transsynaptic tracing experiments^56^ to identify SC neurons that receive inputs from the ACC using the AAV1-CAG-Flpo virus described above. We produced the Flp-dependent tdTomato reporter line Ai65F by crossing the Cre- and Flp-dependent tdTomato double-reporter line Ai65D ^73^ (Jackson Laboratory 021875) to the Cre deleter line Meox2-Cre ^74^ (Jackson Laboratory 003755), so that only Flp is required for expression of tdTomato. An AAV1-CAG-Flpo virus was injected in the ACC of these mice to label postsynaptic neurons with tdTomato. After allowing 4-6 weeks for Flpo and tdTomato expression, we sectioned the brain into 50µm slices and used standard immunohistochemistry techniques to identify GABA-expressing neurons co-labeled with tdTomato. The tissue was placed in 5% normal goat serum, 1% triton blocking solution in 0.1M phosphate buffered saline (PBS) for 1 hour at room temperature. It was then incubated in primary antibodies against GABA and NeuN (rabbit anti-GABA, 1:500, A2052 Sigma; guinea pig anti-NeuN, 1:500, 266-004 Synaptic Systems) and 1% NGS/ 0.5% triton overnight at 4°C. Following washing in 0.1M PBS 3×10’, the tissue was incubated in secondary antibody (goat anti-rabbit IgG AlexaFluor 488, goat anti-guinea pig IgG AlexaFluor 647, 1:500, Invitrogen) and 1% NGS for 4 hours at room temperature. Tissue was then washed 3×10’ in PBS, mounted, and coverslipped with anti-fade mounting medium (Prolong Gold, ThermoFisher). Tiled z-stacks were collected on a confocal microscope (SP8, Leica) using a 20x objective at 1024×1024 resolution, 2 µm apart (∼20 z-slices) from sections containing the SC (between -3.4 and -4.7 AP from bregma). A 1×1 mm ROI was selected over the intermediate and deep layers of SC and z-projected across 25µm. Background was automatically subtracted (ImageJ), and NeuN+ cells were identified using an automated cell-counting binary mask (watershed segmentation, ImageJ). GABA+/NeuN+ and tDTom+/GABA±/NeuN+ cells were manually identified and calculated as the proportion of NeuN+ cells (cell-counter plugin, ImageJ).

### Behavioral apparatus and task training

Mice were trained to report the spatial location of visual cues by rotating a trackball left or right with their forepaws, similar to a previous design^10^. Animals were headfixed on a behavior rig assembled from optical hardware (Thorlabs), placed in a polypropylene tube to limit body movement, and positioned ∼8 cm from an LCD screen (7” diagonal; 700YV, Xenarc Direct) such that their forepaws rested on a trackball. Ball movements were monitored with a commercially available USB optical trackball mouse (Kensington Expert Mouse K64325). The original trackball was replaced with a 55mm diameter ping pong ball (Joola), which was light enough for mice to rotate comfortably. We inserted a hypodermic tube down the center of the trackball so it could only be rotated along a single axis (left or right). To fix the ping pong ball to the optical mouse, we made grooves in the trackball chassis and secured the hypodermic needle with hot glue. A USB host shield (SainSmart) was used to connect the output of the optical mouse to a microcontroller (Arduino), which ran a custom routine that detected ball movements every 10ms. In the event of a ball movement, the microcontroller outputted a timestamp and the amount of movement (in pixels) to a behavioral control computer (Dell). In addition, a time stamp was sent every 100ms to synchronize timing between the microcontroller and the behavior computer. In our system, one pixel of optical mouse movement corresponded to ∼0.15° movement on the ball. Behavioral control was implemented with custom software written in MATLAB (Mathworks) using the Psyhcophysics and the Data Acquisition toolboxes.

During the inward task (Figures 2-5), the presented visual cue (black square, ∼20°) started on either the left or the right side of the LCD screen. The trackball controlled the location of the visual stimulus in closed loop in real-time. The gain of coupling between the trackball and the stimulus was calibrated so that rotating the ball by the threshold amount (15°) in the correct direction moved the stimulus from its starting position to the center of the screen. Ball positions were accumulated throughout the trial until the stimulus reached the center or the response window expired. Under this closed-loop control, any movement in the incorrect direction displaced the stimulus farther away from the center and towards the edge of the screen. Hence, such movements had to be offset by additional movements in the correct direction for the stimulus to move to the center of the screen and for the trial to be considered correct. If the ball was moved opposite to the direction of the instructed cue by the threshold amount, the stimulus moved to the edge of screen and the trial was considered incorrect. The software operated similarly when mice were trained on the outward contingency task (Figure 6), except that they were required to move the visual cue to the edge of the screen for the trial to be considered correct, and moving the stimulus to the center of the screen was considered an incorrect response. Water was given as reward on correct trials, which was delivered through a metal spout placed within the reach of the tongue; the amount dispensed was controlled by opening a solenoid valve for a calibrated period.

The following procedures were used to train mice on both the inward (Figure 2A) and outward (Figure 6A) task contingencies. Mice were taken through successive stages of training until they became proficient at the task. Once mice recovered from the surgery, they were water restricted for 5-7 days (≥ 1ml/day) and then trained to lick a metal spout to obtain small water rewards (3-6 µL). If mice did not receive their water allotment during training, they were given the remaining amount as hydrogel (Clear H2O) in their home cage. After mice reliably licked the waterspout, they earned water rewards by using the trackball to move the presented stimulus to the center of the screen. To discourage spontaneous trackball movements, mice were required to hold the ball still for 1s to trigger trial start, which was signaled with an auditory tone (0.5s, 1 KHz); the visual cue appeared with a 1s delay after the onset of this tone. During early stages of training, only movements in the correct direction contributed to movement of the stimulus. Once mice reliably moved the ball in either direction on >90% of trials, this condition was removed, and the movement of the stimulus was fully coupled to the movement of the trackball. In the next stage of training, we used an anti-bias algorithm in which the same stimulus was repeated on consecutive trials if mice made an error until they performed the trial correctly; stimulus location on trials following correct trials were randomly chosen. Once performance reached ∼70%, the anti-bias algorithm was turned off and stimuli were presented in a randomized manner. Throughout all stages, auditory white noise was used to signal miss trials if mice failed to move the ball to threshold before expiration of the response window. As mice progressed through the training stages, we gradually decreased the response window from 10s to 1s. Correct and incorrect trials were signaled with auditory tones (0.2 kHz and 10kHz, respectively), followed by an inter-trial delay of 2.5s.

Animals trained for two-photon imaging experiments were taken through two additional stages of training. First, we turned off the closed loop coupling between the trackball and the stimulus, and instead flashed stimuli for 200ms. This was to avoid the potential of evoking neural activity due to the movement of the visual cue on the screen. Second, we introduced uncertainty in temporal expectancy for stimulus onset by randomizing the period between the auditory cue signaling trial start and the onset of the visual stimulus (exponential distribution with mean of ∼1.8 seconds, min and max delay of 1 and 5s, respectively). Only mice trained for imaging experiments were taken through these steps.

A cohort of untrained mice were used to test the role of the SC in spontaneous trackball movements. Mice received an injection of AAV-syn-Jaws in the SC and implanted with an optic fiber cannula over the injection site, as described above. After mice recovered from surgery, they were water restricted for 5-7 days and then trained to lick a metal spout to obtain small water rewards (3-6 uL). Once mice reliably licked the waterspout, we began sessions of randomly rewarded no-stimulus trials. During these sessions, no visual or auditory cues were presented that would signal the beginning or end of each trial. No-stimulus “trials” of 2s duration were presented continuously, such that there was no delay between the end of one trial and the beginning of the next (note that this “trial” structure was not apparent to the mice, and is only used for administering rewards, optogenetic stimulation, and subsequent analysis). On each trial, one movement direction, clockwise or counterclockwise, was randomly selected by the software to be rewarded. If mice moved the ball in the designated direction during this period, they obtained a small water reward; no reward or feedback was provided for no-movement or “incorrect” trials (in which mice moved the ball to threshold, but in the unrewarded direction). To encourage movements in both directions, we simultaneously used two anti-bias algorithms during each session: 1) the movement direction selected for rewarding was repeated on consecutive trials until mice received a reward; and 2) the rewarded movement direction switched once mice received a reward.

### Optogenetic manipulation of behavior

Photostimulation was provided with a solid state 593nm laser for experiments using Jaws (OptoEngine). Laser stimulation was triggered from the behavior control computer and lasted from 0.3s before to 1s after visual cue onset. ∼20% of trials were pseudorandomly selected for photostimulation. We additionally imposed the condition that photostimulation could not occur on two consecutive trials. The output of the laser was coupled to a patch cable (Thorlabs) with a FC/PC fiber coupler (OptoEngine). Laser power through the patch cable was measured with a digital power meter before each experiment (Thorlabs). In experiments requiring photostimulation through chronic windows over the visual cortex, the ceramic ferrule of the patch cable was positioned so it filled the entire 3mm window and delivered a constant light pulse with 20mW of power. In experiments requiring light delivery through implanted optical fiber, the ceramic ferrule of the patch cable was coupled to the fiber optic cannula with a ferrule mating sleeve (Thorlabs). Activity in the superior colliculus was inhibited by delivering 10mW of constant yellow light. Photoinhibition of ACC outputs was achieved by constant yellow light (20mW) illumination through an implanted fiber optic cannula (SC) or through a chronic window (VC).

For optogenetic manipulation of spontaneous movements in untrained mice, an average of 39% (±15%) of “trials” were pseudorandomly selected for photostimulation, with the additional stipulation that there be at least six consecutive trials (∼12s) between laser trials. Laser stimulation was triggered from the behavior control computer and lasted 1.3s, beginning from 0.3s before the (uncued) trial start.

### Behavioral analysis for optogenetic experiments

Trials were labeled as ipsi or contra depending on the location of the presented visual cue relative to the brain hemisphere that was photostimulated. We computed several metrics to quantify the performance on each trial type. The timeout rate is the proportion of trials in which the ball was not moved to threshold within the allotted response window out of all trials where a given stimulus was presented. The incorrect rate is the proportion of trials in which the ball was moved opposite the direction instructed by the cue out of all completed trials for each visual stimulus (i.e., excluding timeout trials). In other words, 100 – incorrect rate equals the correct rate. The response time was the amount of time taken from stimulus onset to a complete correct response. The response window (i.e., time given for making a complete response) was set to 1s.

Inactivation-induced change in counterclockwise (CCW) action bias shown in Figure 5D and 6E was calculated as follows for the inward contingency task:

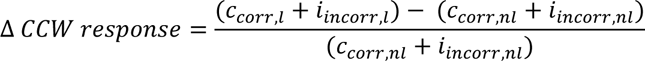

and for the outward contingency task:

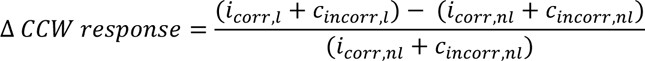

The contra stimulus (CS) action bias in Figure 6D was computed as follows for both the inward and outward tasks:

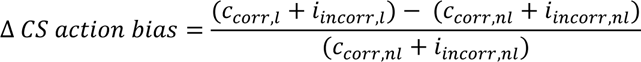

In the equations above, *c* and *i* refer to contra and ipsi cues, *corr* and *incorr* refer to correct and incorrect rates, and *l* and *nl* refer to laser and non-laser trials, respectively.

We quantified the effect of optogenetic inactivation on inward and outward tasks by comparing behavioral performance on non-laser and laser trials from the same sessions. Data from three optogenetic sessions were combined for each mouse. We used a permutation test to quantify the significance of changes in behavioral performance observed with optogenetic inactivation. We tested against the null hypothesis that the observed change in performance does not depend on laser inactivation. In each round of permutation, we randomly reassigned the laser labels across trials for each animal in an experiment. We then concatenated trials for all animals and recalculated the average laser-induced change in the incorrect rate, the timeout rate, and the response time for contra and ipsi cue trials. This process was repeated 1000 times, creating a distribution of performance changes expected by chance. The two-tailed p-value was computed as the proportion of performance changes that were as or more extreme than the observed change on either side of the distribution. We determined significance using the Bonferroni adjusted alpha-value of 0.05/2 = 0.025 for analyses that compared the effect of inactivation on contra and ipsi cue trials.

We also tested the effect of optogenetic inactivation on spontaneous counterclockwise and clockwise movements in untrained mice. Only trials with attempted movements, regardless of whether they were rewarded, were considered for analysis. Trials in which the ball was moved by the threshold amount of 5° were labeled as CW and CCW based on the direction of movement. Inactivation induced changes in CW and CCW movements, shown in Supplementary Figure 3, was calculated as follows:

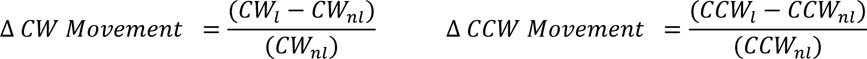

In the equations above, *l* and *nl* refer to laser and non-laser conditions, respectively; CW and CCW refer to clockwise and counterclockwise direction of movements, respectively. We quantified the effect of optogenetic inactivation by comparing attempted movements of each type in laser and non-laser conditions. Though inactivation for this experiment was performed on either the left (n = 2 mice) or right SC (n = 3 mice), movement directions were transposed in the analysis to make all consistent with a left SC inactivation. That is, for mice with right SC inactivation, CW movements were coded as CCW, and CCW movements were coded as CW; movement coding for mice with left SC inactivation were left as is. Importantly, left and right SC inactivation had opposite effects on movements; left SC inactivation decreased counterclockwise actions, whereas right SC inactivation decreased clockwise actions. Trials from 4 sessions were concatenated for each mouse. We used one-sample t-tests to test against the null hypothesis that observed changes in CW or CCW movements for each mouse were significantly different from no change/zero. A two-sample t-test was used to determine if changes in CW movements and CCW movements were significantly different from each other.

### Two-photon microscopy

GCaMP6s fluorescence was imaged through a 16x/0.8 NA objective (Nikon) using galvo-galvo scanning with a Prairie Ultima IV two-photon microscopy system. Excitation light at 910nm was provided by a tunable Ti:Sapphire laser (Mai-Tai eHP, Spectra-Physics) equipped with dispersion compensation (DeepSee, Spectra-Physics). Emitted light was collected with GaAsP photomultiplier tubes (PMTs; Hamamatsu). A blackout curtain attached to a custom stainless-steel plate (eMachineShop.com), which was mounted on the headplate to prevent light from the LCD screen from entering the PMTs. Layer 5 GCaMP6s-expressing neurons in transgenic animals were imaged with 2x optical zoom, 350-550µm below the brain surface for 20-minute-long behavioral sessions. Upon completion, excitation wavelength was changed to 830nm, which produced both a signal from the retrobeads and a structural green signal from GCaMP6. Emission signals were split with a dichroic mirror (FF649-Di01; Semrock) mounted in a filter cube and directed to two different PMTs. Simultaneous imaging of both signals with 830nm excitation facilitated spatial alignment of retrobead signals to GCaMP6s signals collected during behavior and allowed us to identify SC-projecting ACC neurons (see ‘Image analysis’). 4x optical zoom was used for imaging callosal or visual cortex axons in the ACC up to 100µm below the surface. All imaging experiments were performed at a frame rate of 5Hz. Laser power at the specimen was controlled with pockel cells and ranged from 10 to 50 mW, depending on GCaMP6 expression levels and depth.

### Image analysis

Images were acquired using the PrairieView software (Bruker) and saved as multipage TIFF files using ImageJ (NIH). Image processing and region of interest (ROI) selection was performed in ImageJ (NIH). To correct for lateralized movements in the x-y axis, images were realigned to a reference frame (the pixel-wise mean of all frames) using the template matching plugin ^75^. ROIs were drawn manually over visually identified neurons using reference frames generated by taking the pixel-wise maximum, mean, and standard deviation projection of all frames. For each neuron, the same shaped ROI was also placed in an adjacent area devoid of other neurons to estimate the background neuropil signal. To minimize the contribution of the neuropil signal to the somatic signal, corrected neuronal fluorescence time series was estimated as F(t) = F_raw_soma(t)_ – 0.7 × F_raw_neuropil_(t)^76^. Similar image analysis was used for axonal imaging experiments, except ROIs were drawn over visually-identified boutons and fluorescence signals were not adjusted for neuropil contamination. ΔF/F (DFF) for each neuron or bouton was calculated as ΔF/F(t) = 100 × (F(t) – F_0_)/F_0_, where F_0_ was the fluorescence value with the highest density (estimated using MATLAB function ksdensity).

To identify retrobead-containing neurons, a reference frame was generated by taking the pixel-wise mean of realigned GCaMP6s frames acquired during the behavioral session at 910nm. The green channel acquired with excitation at 830nm was realigned to this reference using the template matching plugin. The resulting translation values were then applied to the channel containing signals from retrobeads. An average projection was taken after this realignment and superimposed onto the 910 nm GCaMP reference frame used for drawing ROIs. Neurons containing retrobeads were then visually identified.

### Analysis of visual activity in axons

We assayed the visual responsiveness of visual cortex or callosal axons in the ACC in animals passively viewing visual stimuli presented at the same locations as during the task (solid black square for visual cortex axons, gratings drifting at 90° for callosal axons; 1s stimulus duration). Visually responsive boutons were identified by comparing pre-stimulus activity averaged over a 1s period to averaged activity 0.6s-1.2s after stimulus onset (two-sided Wilcoxon signed-rank test, P < 0.01). AUROC values were computed for each bouton using stimulus activity on contra and ipsi trials. A preference score was then computed as 2 x (AUROC – 0.5); this score ranged from 1 (complete contra preference) to -1 (complete ipsi preference). To determine if individual boutons had significant stimulus preference, scores were recomputed after shuffling trial labels 1000 times. Observed preference scores outside the center 95% of the shuffled distribution were considered significant (P < 0.05, two-sided).

### Analysis of task responses of ACC neurons

For imaging experiments during the inward contingency task (Figures 2, 3), the session-wide DFF trace for each neuron was z-score normalized. Responses on individual trials were aligned to visual stimulus onset. Trials were labeled as contra-counterclockwise, contra-clockwise, ipsi-counterclockwise, and ipsi-clockwise depending on the location of the visual cue relative to the hemisphere imaged and the action selected by the animal. We only analyzed sessions with at least 5 trials in each condition (note that the contra-clockwise and ipsi-counterclockwise conditions correspond to incorrect trials). The color plots in Figure 2F and 3D were generated by averaging responses on each trial condition for each neuron. Note that rows correspond to neurons, and each row plots activity of the same neurons across the four conditions. For analyses in Figures 2G, H and 3E, responses were averaged over a 1s post-stimulus period (post-stimulus response) and then compared for the indicated trial conditions using Wilcoxon signed-rank tests. To determine trial activity on contra cue trials (Figure 2H), we first separately averaged post-stimulus responses on contra-counterclockwise and contra-clockwise trials; responses on these trial types were then averaged together to generate contra responses. Trial activity on ipsi cue, counterclockwise, and clockwise trials was determined similarly, except post-stimulus responses on the following trial types were used, respectively: 1) ipsi-counterclockwise and ipsi-clockwise; 2) contra-counterclockwise and ipsi-counterclockwise; and 3) contra-clockwise and ipsi-clockwise.

### Decoding actions from neuronal activity

We built linear support vector machine (SVM) classifiers using the LIBSVM library for MATLAB^77^ to test whether the activity of ACC-SC neurons can be used to predict which action was selected by the animal. Selection of the regularization parameter C was performed with a dataset not included in the analysis (unlabled ACC neurons). We tested a range of values for C and selected the one which gave the best classifier accuracy on a 10-fold cross-validated dataset. For this procedure, the classifier was trained to distinguish between contra-CCW and ipsi-CW trials.

Since we simultaneously recorded only a few ACC-SC neurons, we constructed ‘pseudotrials’ by combining neuronal responses recorded across different sessions. SVM classifier was trained to predict CCW or CW trials based on post-stimulus activity (z-scored DFF averaged over 1s after stimulus onset) on individual trials. Activity for the CCW label was taken from contra-CCW and ipsi-CCW, and for the CW label from contra-CW and ipsi-CW trials (also see schematic in Figure 3F). We ran 1000 iterations of the model. There was an unequal number of trials between conditions, so we used subsampling to balance trial types on each iteration (5 trials from each condition). We minimized model over-fitting by using the cross-validation technique (10-fold) to split the data into a training and testing set on each iteration. Classifier performance on each iteration was estimated by averaging prediction accuracies across the 10 splits. Final classifier accuracy was determined by averaging these mean accuracies across all iterations. To determine if the action decoder performed above chance, we used an identical procedure except we shuffled labels for the test data. Mean prediction accuracy derived from correctly labeled test data and falling outside the center 95% of the shuffled distribution was considered significant.

### Electrophysiological recordings in the SC

For photostimulation of the ACC, we injected AAV1.CaMKII.hChR2(H134R)-mCherry (University of Pennsylvania vector core) at coordinates AP: 0.5mm, ML: 0.5mm, and DV: 0.4 and 0.9mm (250 nL at each site). Cannulas were implanted such that the optical fiber was 0.3mm below the pia (300µm/0.39NA core fiber optic coupled to a 2.5mm stainless steel ferrule; CFMC13L02, Thorlabs). We tested the effect of photostimulating ChR2-expressing ACC neurons on activity in the SC using a 473nm blue solid state laser (Optoengine). One or two days before the experiments, mice were habituated to head-fixation in 1h sessions. On the day of the experiment, mice were anesthetized with isoflurane and the Metabond and silicone elastomer on the skull were removed. The mouse was placed on the stereotaxic frame and a 500 μm diameter craniotomy was performed on top of the recording site (from bregma: -3.6 to -4 mm anteroposterior and 0.8 to 1 mm mediolateral). The dura above the cortex was removed and the craniotomy was protected with saline and a piece of gelfoam. SC craniotomy was performed on the same side as that implanted with the fiber optic cannula over the ACC. The skull was covered again with silicone elastomer and the animal was returned to its home cage to recover from anesthesia for at least 2 hours. After recovery, mice were headfixed and the silicone and gelfoam overlaying the craniotomy was gently removed. 0.9% NaCl solution was used to keep the surface of the brain wet for the duration of the recordings.

After placing the animal in the recording set up, we submerged a reference silver wire in the NaCl solution on the skull surface. The position of the 16-channel silicone probe (A1x16-Poly2-5mm-50s-177-A16, NeuroNexus) was referenced on lambda and the surface of the brain and lowered slowly (1 min per mm) to reach superficial sensory layer of the superior colliculus (∼1.3 mm in the ventral axis) using a motorized micromanipulator (MP – 285; Sutter Instrument Company). The extracellular signal was amplified using a 1x gain headstage (model E2a; Plexon) connected to a 50x preamp (PBX-247; Plexon) and digitalized at 50 kHz. The signal was highpass filtered at 300Hz. Once the visual layer of the SC was identified (characterized by strong and reliable visual responses to drifting gratings and sparse noise), the recording probe was lowered ∼400 μm deeper to the motor layer of the SC. For successful recordings, the silicone probe was gently retracted and the recording tract was marked by re-entering the DiI coated probe (2 mg/mL – D3911, ThermoFisher Scientific) at the same location. For some experiments, we were able to record from two locations spaced 500μm apart in the dorsal-ventral axis. The brain was harvested post-hoc and sectioned to confirm the probe location and ChR2 expression in the ACC. Spikes were isolated online with amplitude threshold using Plexon Recorder software, but re-sorted using the MountainSort automated spike sorting algorithm^78^. Units were curated manually after automatic detection.

Visual stimuli were presented during recordings in the SC to increase neuronal responsiveness. Sparse noise on a 3 × 5 grid (square size was the same as used for behavior experiments) of black and white square on a gray background (50% luminance) were displayed for 0.1s, followed by a 0.1s gray screen period. Positions were randomized within each block such that black and white squares were presented once at each of the 15 positions. The total duration of a block was 6 sec, with 1s inter-block intervals. Photostimulation of the ACC (10ms blue light pulses at 20 Hz) was performed on 50% of the blocks. Photostimulation started 0.5s before visual stimulus presentation and was turned off 0.5s after stimulus offset (total duration of 7s).

Since ACC axons in the SC target the intermediate and deep layers, we focused our analysis on recordings made from these areas. While we observed robust retinotopically organized visual responses in the superficial layer, we rarely encountered cells in the deeper layers that specifically responded to the location of sparse noise stimuli. Therefore, recordings from deeper neurons likely reflect ongoing, spontaneous activity that is modulated by the visual stimulation. We first determined if individual neurons were significantly modulated by laser activation of the ACC by comparing the firing rates (FR) of activity on non-laser and laser trials. For each modulated neuron, we also computed a laser modulation index using the following equation:

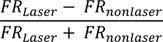

## Notes

### Competing Interest Statement

The authors have declared no competing interest.

